# Structural mechanism of insulin receptor activation by a dimeric aptamer agonist

**DOI:** 10.1101/2025.05.12.653381

**Authors:** Junhong Kim, Hyeonjin Na, Si-Young Choi, Eun Ju Oh, Hyunsook Lee, Sung Ho Ryu, Na-Oh Yunn, Yunje Cho

**Author notes:** These authors contributed equally to this work (Junhong Kim, Hyeonjin Na).

## Abstract

Insulin binding to the insulin receptor (IR) triggers signaling pathways that regulate glucose uptake and cell growth. In previous work, we identified a DNA aptamer, A62, which partially activates the IR. During engineering aptamers for improved *in vivo* stability, we discovered that crosslinking two A62 aptamers with linkers of varying lengths led to full phosphorylation of the IR, though activation remained selective to the AKT pathway. To elucidate the mechanism behind this aptamer-induced full activation of the IR, we determined the structure of the IR in complex with a dimeric form of A62 (A62D) linked by an 8-nucleotide connector. We identified three distinct conformations of the IR: arrowhead-shaped, pseudo-arrowhead-shaped, and pseudo-gamma-shaped. The pseudo-gamma-shaped conformation closely resembles the structure of a fully active IR bound by a single insulin molecule. In these configurations, only one A62 monomer (A62M) within the A62D dimer binds to the IR dimer. This binding brings the IR monomers into close proximity, promoting intermolecular trans-phosphorylation. Our findings provide valuable structural insights for the development of novel therapeutic strategies targeting the IR.

## Introduction

Insulin is a peptide hormone produced by the beta cells of the pancreatic islets. Upon binding to the insulin receptor (IR) in peripheral tissues, insulin activates the receptor, triggering intracellular signaling cascades through the phosphoinositide 3-kinase (PI3K)/AKT and mitogen-activated protein kinase (MAPK) pathways (1–3). Therefore, insulin-mediated IR activation plays a pivotal role in regulating glucose homeostasis, protein synthesis, lipogenesis, cell growth, proliferation, and development (4, 5). Dysregulation of IR signaling leads to chronically elevated blood glucose levels, resulting in diabetes mellitus (6). Individuals with diabetes are prone to a wide range of complications and have an increased risk of cancer (7).

The IR is a homodimer, with each protomer consisting of an α chain and a β chain (8). The α-subunit and N-terminal portion of the β-subunit are extracellular, whereas the β-subunit spans the membrane and contains a cytoplasmic tyrosine kinase domain responsible for insulin-dependent phosphorylation. The α-subunit mediates ligand binding via a leucine-rich repeat (L1), a fibronectin type III domain (FnIII-1), and an α-helical C-terminal domain (αCT). Biochemical and structural studies of insulin binding to IR suggest that each IR protomer contains two insulin-binding sites—primary site (site-1) and secondary site (site-2) (9–13). Site-1 is formed by the L1 and αCT′ domains (with the prime indicating structural elements from the opposite protomer), and interactions at this site are crucial for insulin binding and IR activation (12, 13). Site-2 comprises residues on the surface of the FnIII-1 β-sheet (9–11).

Structural studies have revealed various conformations of the extracellular domain of IR in both the apo state and when bound to different numbers of insulin molecules (9–24). In the apo state, the two αβ monomers form a symmetric, Λ-shaped dimer in an antiparallel orientation, with multiple interfaces between protomers (17, 18). A single insulin molecule binds simultaneously to the L1 and αCT′ domains at site-1 of the IR, resulting in an asymmetric gamma (Γ)-shaped structure (12, 13, 21, 24, 25). In this Γ-shaped structure, the L1, CR, L2, and FnIII-1 domains in one protomer form an upshifted “head,” while the corresponding domains in the opposite protomer form a downshifted head. The stalks are composed of the FnIII-1, FnIII-2, and FnIII-3 domains in both protomers. Upon insulin binding, the L1 and αCT′ domains at site-1 are lifted toward FnIII-1′ relative to their position in the apo state. When multiple insulin molecules bind, the IR adopts a series of tilted T-shaped structures, reaching a fully T-shaped conformation when all four insulin-binding sites are occupied (9–11, 18, 22–24). Insulin binds to IR with negative cooperativity, and under physiological conditions, the binding of a single insulin molecule is sufficient to fully activate the receptor (26, 27).

Although insulin has been the most commonly issued treatment for diabetes, considerable effort has been directed toward developing alternative therapeutics. These include peptides that selectively activate the AKT signaling pathway (20–22, 28), antibodies that enhance IR activation in the presence of insulin (29, 30), and DNA aptamers (24, 31, 32). In a previous study, we developed the A62 aptamer, a selective partial agonist for IR (31). The A62 aptamer induces mono-phosphorylation of the receptor by binding at the interface between the L1 and FnIII-1’ domains of the opposite IR protomer (24). Two A62 aptamers bind symmetrically to the IR dimer, inducing an arrowhead-shaped conformation. Additionally, a single A62 aptamer can bind IR together with insulin. In both cases, the membrane-proximal ends of the FnIII-3 domains are separated by ∼40 Å, which may explain the aptamer’s ability to induce only mono-phosphorylation of the IR.

One major obstacle to the clinical use of aptamers is their small molecular size, which can limit their stability *in vivo*. To address this issue, we aimed to enhance the *in vivo* stability of the A62 aptamer agonist for IR by creating dimeric forms. We crosslinked two A62 molecules using linkers of various lengths. The resulting A62 dimers, in which two A62 monomers (A62M) were connected by linkers ranging from 8 to 19 nucleotides, induced full phosphorylation of the IR. To understand the basis of this aptamer-induced full phosphorylation, we determined the structure of the IR bound to an A62 dimer linked by 8 thymine nucleotides (A62D-8T) using cryogenic-electron microscopy (cryo-EM). Structural analysis revealed three distinct IR conformations: arrowhead-shaped, pseudo-arrowhead-shaped, and pseudo-gamma-shaped, which we refer to as IR_arrowhead_, IR_pseudo-arrowhead_, and IR_pseudo-gamma_, respectively. Unlike the symmetric IR_arrowhead_ dimer, IR_pseudo-arrowhead_ and IR_pseudo-gamma_ form asymmetric dimers. In these configurations, each A62M module of the A62D binds to the IR dimer, bringing the IR protomers into close proximity to facilitate trans-phosphorylation. Our findings suggest a strategy for inducing full IR phosphorylation by crosslinking aptamer modules that bind to the IR dimer, thereby promoting intermolecular trans-phosphorylation.

## Materials and Methods

### Reagents and antibodies

The aptamers used in this study were synthesized by Aptamer Science, Inc. (Pohang, Korea). The anti-insulin receptor β-subunit (CT-3) antibody was purchased from Santa Cruz Biotechnology (Santa Cruz, CA, USA). Antibodies specific for phosphorylated insulin receptor at tyrosine residues Y1146, Y1150, and anti-phospho-tyrosine (4G10) were purchased from Millipore. Additional antibodies against phosphorylated insulin receptor at residues Y1322, Y1316, Y1150/Y1151, and Y960 were purchased from Invitrogen (Carlsbad, CA, USA). The anti-phospho-ERK1/2 (T202/Y204), anti-phospho-AKT (T308), and anti-phospho-AKT (S473) antibodies were purchased from Cell Signaling Technology (Danvers, MA, USA). Secondary antibodies, goat anti-rabbit IgG and anti-mouse IgG conjugated to DyLight 800, were also purchased from Invitrogen.

### IR phosphorylation assay

Rat-1 cells overexpressing human IR (Rat-1/hIR) were cultured in high-glucose Dulbecco’s modified Eagle medium (DMEM) supplemented with 10% (v/v) fetal bovine serum (FBS; Gibco) and antibiotic-antimycotic solution (Gibco). All cells were incubated at 37°C in a humidified atmosphere with 5% CO_2_ until used for experiments. Aptamers and insulin were prepared in Krebs–Ringer HEPES buffer (25 mM HEPES, pH 7.4) containing 120 mM NaCl, 5 mM KCl, 1.2 mM MgSO_4_, 1.3 mM CaCl_2_, and 1.3 mM KH_2_PO_4_. To reconstitute the tertiary structure, all aptamer samples were heated at 95°C for 5 min and then slowly cooled to room temperature. Before stimulation with insulin or aptamers, cells were serum-starved for 3 h. They were then treated with the indicated concentrations of insulin or aptamers for the specified duration. After stimulation, the cells were washed with cold phosphate-buffered saline and lysed in a cell lysis buffer (50 mM Tris-HCl, pH 7.4) containing 150 mM NaCl, 1 mM EDTA, 20 mM NaF, 10 mM glycerophosphate, 2 mM Na_3_VO_4_, 1 mM phenylmethylsulfonyl fluoride (PMSF), 10% glycerol, 1% Triton X-100, 0.1% SDS, 0.5% sodium deoxycholate, and a protease inhibitor cocktail. The lysates were clarified by centrifugation at 14,000 rpm for 15 min at 4°C, and the supernatant was mixed with 5× Laemmli sample buffer. Proteins were separated by SDS-PAGE, transferred to a nitrocellulose membrane, and blocked with 5% skim milk for 30 min. The membrane was then probed with the appropriate primary antibody overnight at 4°C. Protein detection was performed using the Odyssey infrared imaging system (LI-COR, Lincoln, NE, USA).

### Expression and purification of IR and A62D complex

To express apo-IR, stable cells were cultured in suspension in Freestyle293 medium at 37°C with 8% CO_2_. When the cell density reached ∼4.0 × 10^6^ cells/mL, cells were harvested by centrifugation and resuspended in a buffer containing 20 mM HEPES (pH 7.5), 400 mM NaCl, and 5% glycerol. The cells were lysed on ice using a Dounce Homogenizer (Kimble), followed by solubilization in a buffer consisting of 20 mM HEPES (pH 7.5), 400 mM NaCl, 1% (w/v) n-dodecyl β-D-maltoside (DDM; Anatrace), 0.1% (w/v) cholesterol hemisuccinate (CHS; Sigma), 5% glycerol, and a protease inhibitor cocktail (Roche). Solubilization was performed for 2 h on ice. Solubilized membranes were then isolated via ultracentrifugation using a Ti45 rotor (Beckman) at 130,000 × *g* for 1 h at 4°C.

The supernatant was collected and incubated with anti-Flag affinity G1 resin (GenScript) for 2 h at 4°C. The resin was batch-washed using a washing buffer containing 20 mM HEPES (pH 7.5), 400 mM NaCl, 5% glycerol, 0.1% DDM, and 0.01% CHS. Proteins were eluted with a buffer composed of 20 mM HEPES (pH 7.5), 400 mM NaCl, 5% glycerol, 0.03% DDM, 0.003% CHS, and 0.4 mg/mL Flag peptide. To remove the Flag peptide, eluted proteins were concentrated using an Amicon Ultra centrifugal device (100 kDa cut-off; Millipore) and diluted in a buffer containing 20 mM HEPES (pH 7.5), 105 mM NaCl, 5% glycerol, 0.03% DDM, 0.003% CHS, 5 mM KCl, and 5 mM MgCl_2_. The proteins were concentrated again using an Amicon Ultra device (100 kDa cut-off; Millipore).

To form the complex with the aptamer, the proteins and pre-activated A62D aptamer were mixed at a 1:1 molar ratio and incubated for 1 h on ice. The mixture was then injected onto a Superose 6 10/300 column pre-equilibrated with a buffer containing 20 mM HEPES (pH 7.5), 105 mM NaCl, 5 mM KCl, 5 mM MgCl_2_, 0.03% DDM, and 0.003% CHS. Eluted fractions were pooled and concentrated to 8 mg/mL using a Vivaspin device (100 kDa cut-off; GE Healthcare) for subsequent cryo-EM analysis.

### Cryo-EM sample preparation and data collection

Cryo-EM grids were prepared by applying 3 μl of the sample to glow-discharged holey carbon grids (C-flat 1.2/1.3 Au, 400-mesh; EMS). The grids were then plunge-frozen in liquid ethane using a Vitrobot Mark IV (Thermo Fisher Scientific) with a blot force of 4 for 5 s at 100% humidity and 4°C. Data collection was performed on a Titan Krios G4 microscope operated at 300 kV, equipped with a Gatan K3 summit direct electron detector in fast mode at a nominal magnification of 79,000×. A total of 9,096 micrographs were acquired. The dataset was collected with a pixel size of 1.0902 Å and a defocus range of −1.0 to −2.0 μm. Each micrograph was dose-fractionated over 50 frames, with a total accumulated dose of 50 electrons/Å^2^.

### Data processing for the IR_arrowhead_ structure

Dose-fractionated image stacks from 9,096 movies were imported into CryoSPARC v4.5.3 (33). The images underwent dose-weighting using patch motion correction, followed by contrast transfer function (CTF) parameter calculation via patch CTF estimation. Low-resolution micrographs were excluded, resulting in 8,664 micrographs being selected for further data processing. A total of 1,312,592 particles were extracted using a template picker. After multiple rounds of 2D classification, 38,995 particles were used for *ab initio* reconstruction to generate an initial 3D model. Heterogeneous refinement identified 11,658 particles from one class with an arrowhead shape, which were subsequently used to train Topaz (34).

The trained model was then applied to extract 626,038 particles. Following 2D classification, 48,970 particles exhibiting an arrowhead shape were subjected to homogeneous refinement and non-uniform refinement, and local refinement. This processing yielded a map with a global resolution of 5.02 Å, based on a Fourier shell correlation (FSC) criterion of 0.143.

### Data processing for the IR_pseudo-arrowhead_ structure

To process the pseudo-arrow conformation, we used the same dataset as for the arrowhead shape. A total of 1,421,993 particles were extracted using a template picker. After multiple rounds of 2D classification, 147,154 particles were used for *ab initio* reconstruction to generate an initial 3D model. Heterogeneous refinement identified 15,052 particles from one class with an asymmetric shape, which were subsequently used to train Topaz (34).

The trained model was then applied to extract 1,886,306 particles. Following *ab initio* reconstruction and heterogeneous refinement, 303,989 particles exhibiting a pseudo-arrowhead shape were subjected to additional particle sorting. After two rounds of *ab initio* reconstruction and heterogeneous refinement, two distinct pseudo-arrowhead maps were obtained, consisting of 41,455 and 33,653 particles, respectively. Although the training focused on particles with asymmetric shapes, some particles still adopted an arrowhead shape. Among the 346,320 particles with an arrowhead configuration, further sorting was conducted to isolate those with symmetrical shapes. After several rounds of particle sorting, 18,306 particles were selected for the final 3D sorting process.

The three particle sets (41,455 and 33,653 from the asymmetric class, and 18,306 from the arrowhead shape) were combined for final *ab initio* and heterogeneous refinement. After excluding 6,559 particles corresponding to junk data, a total of 86,854 particles underwent *ab initio*, non-uniform refinement, and local refinement of the extracellular domain. This processing yielded a map with a global resolution of 6.26 Å, based on a Fourier shell correlation (FSC) criterion of 0.143.

### Data processing for the IR_pseudo-gamma_ structure

To process the pseudo-gamma conformation, we used the same dataset as for the arrowhead shape. Using a template picker, we extracted 3,495,704 particles. Following several rounds of 2D classification, 356,821 particles were selected for *ab initio* reconstruction to generate an initial 3D model. After heterogeneous refinement, 24,688 particles belonging to one class with an asymmetric shape were subjected to Topaz training (34). From this trained model, 759,607 particles were extracted. Subsequent *ab initio* reconstruction and heterogeneous refinement resulted in 100,204 particles exhibiting the pseudo-gamma shape, which were then subjected to additional particle sorting. After three rounds of *ab initio* reconstruction and heterogeneous refinement, 42,469 pseudo-gamma particles were identified. Further filtering via 2D classification removed non-relevant particles, leaving 37,078 particles. Ultimately, we achieved a map with a global resolution of 9.35 Å, determined using a FSC criterion of 0.143.

### Model building

The atomic model building of the IR_arrowhead_ conformation began with the docking of the human IR bound to A62M (PDB: 7YQ6) into the map, utilizing UCSF Chimera v1.17.1 (24, 35). Manual adjustments to the model were performed in Coot (36). Since the electron density map of A62D covered only the monomer, we fitted the A62M model accordingly. The model underwent real-space refinement in PHENIX 1.14, applying rigid body and secondary structure restraints (37). The final validated model achieved a MolProbity score of 2.29 with a clash score of 17.7 (38).

For the IR_pseudo-arrowhead_ structures, model building began with the docking of the human IR bound to A62M (PDB: 7YQ6) into the map, utilizing UCSF Chimera v1.17.1 (24, 35). Manual adjustments to the model were performed in Coot (36). Since the electron density map of A62D covered only the monomer, we fitted the A62M model accordingly. The model underwent real-space refinement in PHENIX 1.14, applying rigid body and secondary structure restraints (37). The final validated model achieved a MolProbity score of 2.32 with a clash score of 21.47 (38).

For the IR_pseudo-gamma_ structures, model building was performed by docking the IR bound to A62M and the IR_A43+Ins_ structure (PDB: 7YQ6, 7YQ3) (24). The models were manually refined in Coot (36) and then subjected to real-space refinement in PHENIX 1.14 (37), using rigid body and secondary structure restraints. The refined models achieved a MolProbity score of 2.29 and a clash score of 21.76 (38).

### Oligomerization analysis of IR by size exclusion chromatography

To verify whether the A62D aptamer induces intermolecular interactions between IR dimers, we divided purified IR protein into two fractions. Each fraction was incubated with either the A62M or A62D aptamer at a molar ratio of 1:2 and 1:1, respectively, for 1 h on ice. The mixtures were then injected into a Superose 6 10/300 column pre-equilibrated with buffer containing 20 mM HEPES (pH 7.5), 105 mM NaCl, 5 mM KCl, 5 mM MgCl_2_, 0.03% DDM, and 0.003% CHS. Peaks corresponding to the void volume and the A62M:IR dimer (2:1) or A62D:IR dimer (1:1) complexes were collected. The samples were subsequently concentrated to 0.04 mg/mL using a Vivaspin device (100 kDa cut-off; GE Healthcare) in preparation for negative-stain electron microscopy (EM).

### Negative-stain EM analysis

To prepare negative-stain EM grids, 3.5 μl of the sample was applied to glow-discharged holey carbon grids (Formvar/Carbon, Cu 400-mesh; EMS). The sample was allowed to adsorb onto the grid for 30 s, after which excess liquid was blotted off with filter paper. The grid was then washed twice with deionized water and stained with uranyl acetate solution for 20 s. Following another blotting step, the grid was allowed to dry overnight at 18°C. Images were acquired using a BIO TEM JEM-1011 instrument, operated at an acceleration voltage of 80 kV, and equipped with a Gatan ES1000W CCD camera at a magnification of 200,000x.

### Immunofluorescence Assay

Rat-1/hIR cells were cultured in Dulbecco’s Modified Eagle Medium (DMEM, Serena) supplemented with 10% fetal bovine serum (FBS, Serena), 100 U/mL penicillin, and 100 µg/mL streptomycin. Cultures were maintained at 37°C in a humidified incubator with 5% CO_2_. Before the immunofluorescence assay, cells were seeded on coverslips coated with poly-L-lysine (10 µg/mL). Serum starvation was performed using FBS-free medium for 2 h prior to aptamer treatment. Each aptamer was applied at the indicated concentrations for 1 h. Following aptamer treatment, cells were subjected to an immunofluorescence assay as described in previous studies (39–42). Briefly, cells were fixed with 4% formaldehyde, prepared by diluting a 16% methanol-free formaldehyde solution (Thermo Scientific), and then permeabilized with Tris-buffered saline (TBS) containing 0.5% Triton X-100. The cells were subsequently incubated overnight with a primary antibody targeting the insulin receptor β subunit (Cell Signaling, #3025S) in an antibody dilution buffer (TBS containing 0.1% Triton X-100, 0.1% sodium azide, and 2% BSA). After primary antibody incubation, cells were treated with a secondary antibody (Goat anti-Rabbit IgG (H+L) Cross-Adsorbed Secondary Antibody, Alexa Fluor™ 488, Thermo Fisher) and Phalloidin (Alexa Fluor™11

568 Phalloidin, Invitrogen). Finally, cells were mounted using Vectashield antifade mounting medium (Vector Laboratories).

### Structured Illumination Microscopy (SIM) and Data Analysis

Super-resolution imaging was performed using an AP DeltaVision OMX Ultra High-Resolution Fluorescence Microscope. 3D-SIM was utilized to acquire images with a resolution below the diffraction limit (43). The system employed a 60× oil-immersion objective lens with a numerical aperture of 1.42 and a refractive index of 1.518 (Olympus). Structured illumination (SI) images were processed for alignment, reconstruction, and deconvolution using the SoftWorx software. DAPI, IR, and Phalloidin signals were detected using 405, 477, and 568 nm excitation lasers, respectively. Optical sections were obtained at 125 nm spacing. Representative images presented are maximum intensity projections of the acquired images for each sample. Maximum projection was generated using ImageJ (National Institutes of Health).

For IR puncta analysis, particle analysis in ImageJ was performed. Particles ranging in size from 0.5 µm² to 4 µm² to were analyzed to exclude non-specific signals from the dataset. All graphs and statistical analyses were generated using GraphPad Prism 5 software. P-values were calculated using Student’s t-test, and the results are expressed as mean ± standard error of the mean (SEM), unless otherwise indicated.

## Results

### Design of a full agonist aptamer for IR

In our previous study, we developed IR-A62 (referred to here as A62M) which is a partial agonist that specifically phosphorylates the Tyr1150 residue of IR, and a single administration of A62M temporarily reduced blood glucose levels in diabetic mice (31). However, to assess A62M’s potential in diabetes, its effects following repeated administration over an extended period need to be evaluated. One of the major challenges in developing aptamers for clinical use is their small size. A62M has a molecular weight of ∼9.1 kDa, which is insufficient to avoid rapid clearance from the bloodstream via renal excretion.

A common strategy to improve the pharmacokinetic properties of aptamers is to create multimeric forms (44). However, multivalency can sometimes hinder aptamer activity by causing steric clashes with the target (45, 46). To address this, we examined the activity of a dimeric form of IR-A62. We constructed a series of dimers by linking two A62M aptamers with thymine nucleotide linkers of varying lengths and assessed their ability to activate the IR (Fig. 1a–c). These linkers consisted of 4T, 6T, 8T, 10T, 12T, 14T, 19T, and 24T thymine nucleotides. They are in various range of lengths, from 4T linker with ∼2.5 nm that connect two A62 aptamers in close proximity to 24T linker with ∼15.2 nm that are approximately three times the diameter of the A62 aptamer, ∼5 nm. We used 400 nM A62M and 200 nM A62D-nTs (where “n” represents the number of thymine nucleotides) to compare their activities, as each A62D-nT aptamer contains two functional ligands. Surprisingly, A62 dimers linked by 8T to 19T induced dual phosphorylation at the Tyr1150 and Tyr1151 residues and significantly enhanced AKT signaling (Fig. 1b). In contrast, dimers linked by 4T, 6T, or 24T showed reduced and limited phosphorylation of the IR. Thus, we conclude that A62 dimers linked by a moderately sized linker can induce full phosphorylation of the IR, whereas linkers that are too short or too long impair this ability.

**Figure 1.**
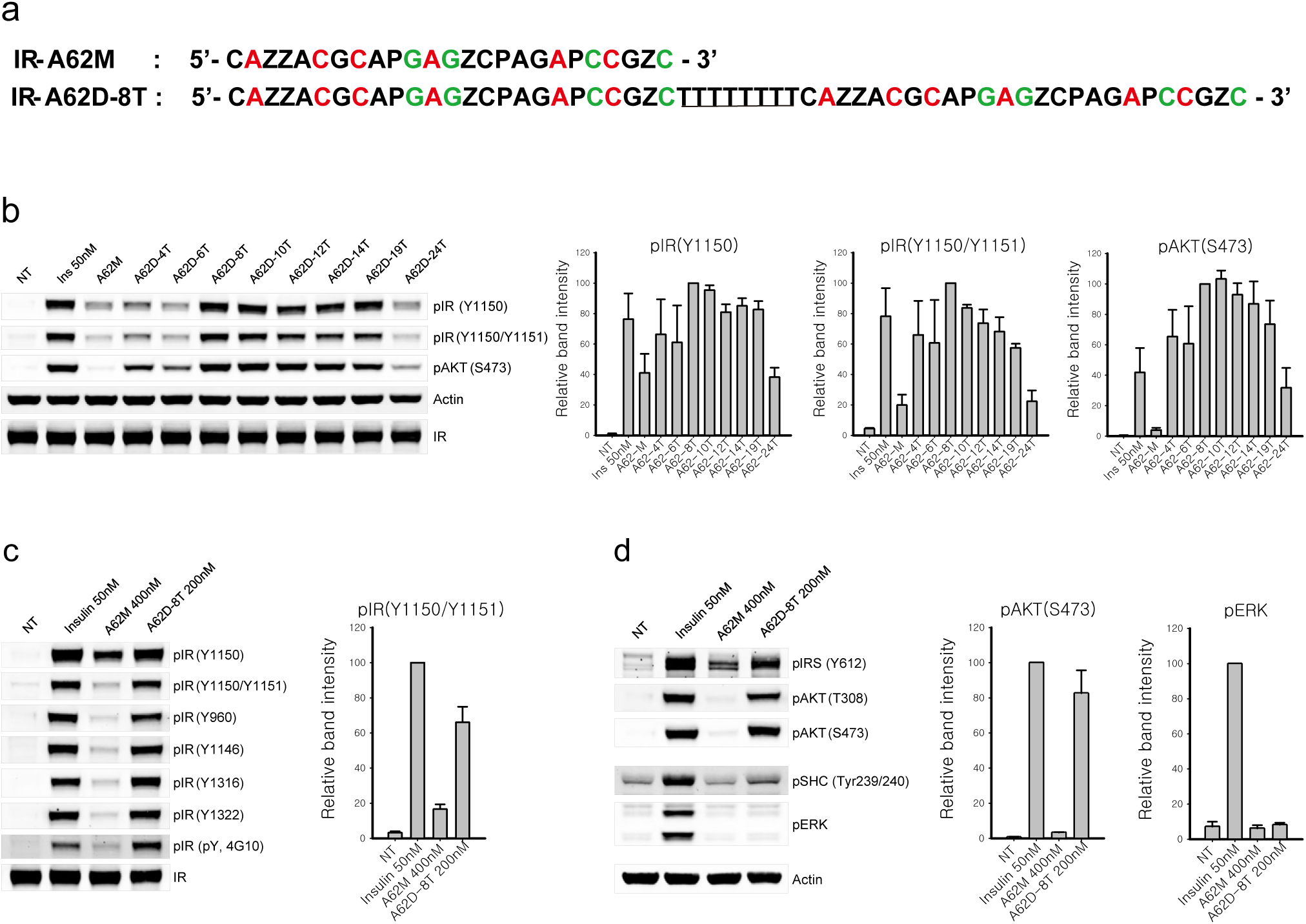
Sequence and agonistic effect on insulin receptor phosphorylation of IR-A62 monomer and IR-A62 dimers. (a) Sequence comparison between the IR-A62 monomer and IR-A62 dimers linked by the 8T linker. For clarity, only IR-A62D-8T is shown. Z and P represent the Benzyl modified- and Naphthyl modified-dU, respectively. Red and Green colored nucleotides represent 2’-fluoro ribose and 2’-O-methyl ribose, respectively. The linker between the two A62M aptamers is underlined. (b) Activity comparison of the IR-A62 monomer and IR-A62 dimers. Rat-1/hIR cells were stimulated with 50 nM insulin for 10 min, 400 nM A62M for 1 h, or 200 nM A62 dimers for 1 h. Bar graphs are presented as the mean ± S.D. of three independent replicates, and normalized by A62D-8T stimulated samples. (c) Phosphorylation of IR and (d) downstream signaling were analyzed using specific phospho-antibodies. Rat-1/hIR cells were stimulated with 50 nM insulin for 10 min, 400 nM A62M for 1 h, or 200 nM A62M-8T. Bar graphs are presented as the mean ± S.D. of three independent replicates, and normalized by 50nM insulin-stimulated samples.

To further characterize the A62 dimer, we selected A62D-8T and measured its effects on the phosphorylation of IR, AKT, and ERK after one hour of stimulation, and compared its activity to A62M. A62D-8T was found to induce phosphorylation of all tyrosine residues, similar to insulin (Fig. 1c). Despite the full phosphorylation of IR by A62D-8T, activation of the MAPK pathway remained significantly lower than that induced by insulin (Fig. 1d).

### Dose and time characterization of the A62D-8T aptamer

We next evaluated the efficacy and potency of A62D-8T in comparison to A62M (Fig. 2a–f, Supplementary Fig. 1). A62D-8T demonstrated significantly higher efficacy in promoting IR and AKT phosphorylation compared to A62M. Specifically, A62D-8T showed greater potency and efficacy in inducing mono- (Y1150), dual- (Y1150/Y1151), and total Tyr (4G10) phosphorylation of the IR. Consistent with previous findings (Fig. 1a–d), A62D-8T increased AKT phosphorylation in a dose-dependent manner, surpassing A62M. However, even at saturation, A62D-8T did not significantly affect ERK phosphorylation.

**Figure 2.**
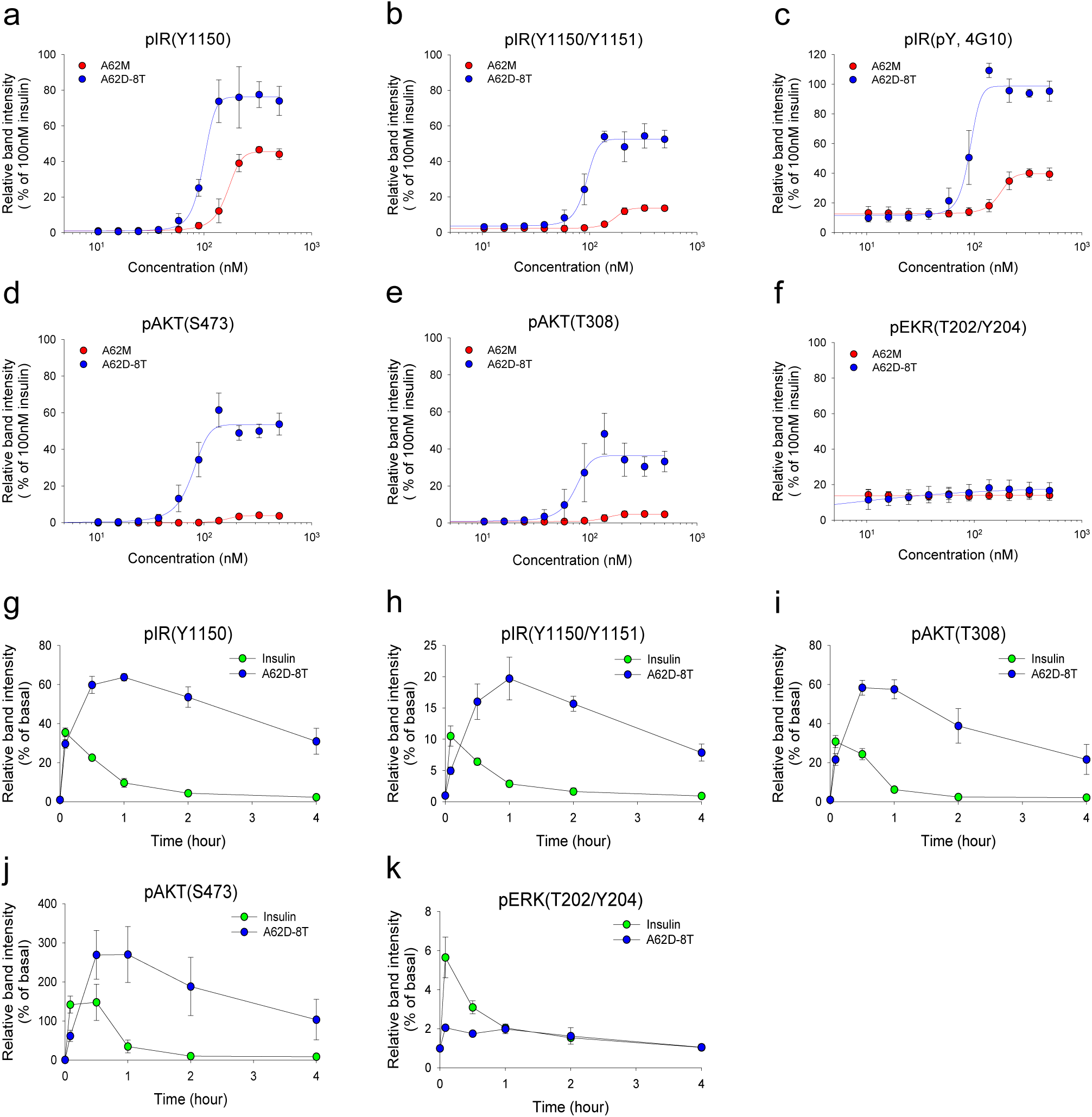
Effects of the dose and duration of A62M and A62D-8T on IR phosphorylation and downstream signaling. Comparison of dose-dependent phosphorylation of IR and downstream signaling by A62M or A62D-8T. Rat-1/hIR cells were incubated with varying concentrations of A62M or A62D for 1 h. The relative band intensities, compared to 100 nM insulin, are shown for (a) pIR Y1150, (b) pIR Y1150/Y1151, (c) pIR pY, (d) pAKT S473, (e) pAKT T308 and (f) pERK T202/Y204, presented as the mean ± S.D. of three independent replicates. Time-dependent phosphorylation of IR and downstream signaling. Rat-1/hIR cells were stimulated with 50 nM insulin for 10 min or 200 nM A62M-8T for 5 min to 4 h. The kinetics of (g) pIR Y1150, (h) pIR Y1150/Y1151, (i) pAKT T308, (j) pAKT S473, and (k) pERK T202/Y204 are presented as the mean ± S.D. of three independent experiments, normalized to each negative control (NT) to determine the fold-change from basal levels.

A notable property of A62M, which differentiates it from insulin, is its slower onset of IR phosphorylation, taking up to 2 h to increase and persisting for over 4 h. Therefore, we also investigated the time-dependent effects of both insulin and A62D-8T (Fig. 2g–k, Supplementary Fig. 1b). Similar to A62M, A62D-8T exhibited a slow and sustained activation of the receptor. Although A62D-8T induced up to a two-fold increase in IR and AKT phosphorylation compared to insulin, it did not significantly alter ERK phosphorylation (Fig. 2a–f). These findings indicate that while A62D-8T fully induces Tyr phosphorylation of the IR, it continues to selectively activate the AKT pathway without triggering the MAPK pathway.

### Structures of the A62D-8T and IR complex

Previously, two A62M aptamers were demonstrated to bind to the IR dimer, inducing an arrowhead-shaped conformation. This structural shift from the apo state resulted in mono-phosphorylation of the IR (24, 31). To explore how A62D-8T induces full phosphorylation of the IR, we determined the structure of the IR bound to A62D-8T. Full-length IR was purified and incubated with A62D-8T at a 1:1 molar ratio. After further purification via size-exclusion chromatography, we resolved the complex structure using cryo-EM (Supplementary Fig. 2–5). Our analysis revealed three distinct conformations of the A62D-8T-IR complex: arrowhead-shaped (IR_arrowhead_), pseudo-arrowhead-shaped (IR_pseudo-arrowhead_), and pseudo-gamma-shaped (IR_pseudo-gamma_) (Fig. 3a–i, Supplementary Fig. 2–5). These conformations were distributed in a ratio of 0.29:0.30:0.12, with resolutions ranging from 6.26 to 9.35 Å.

**Figure 3.**
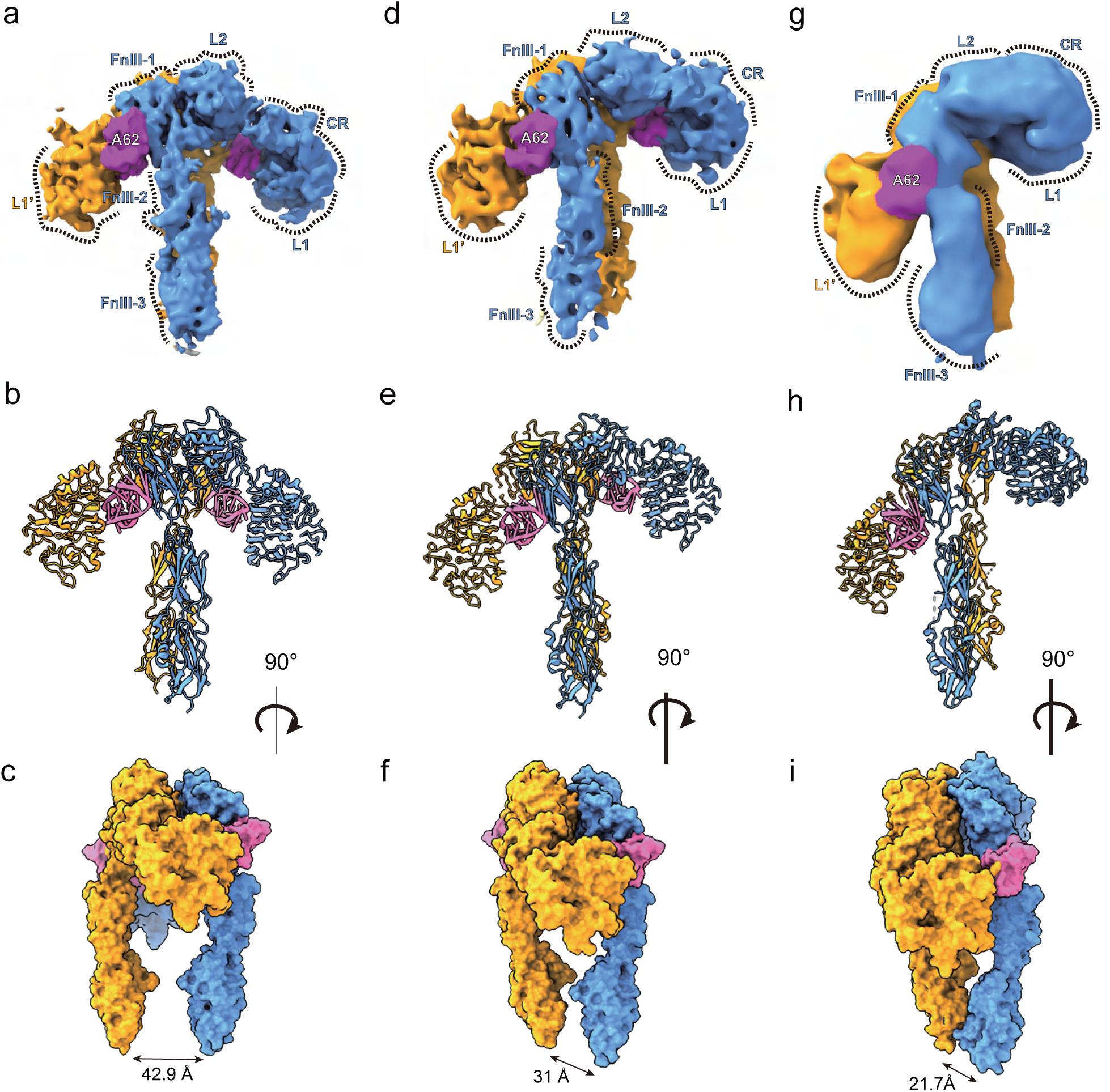
Three distinct conformations of the A62D-8T-bound IR dimer. (a) Cryo-EM map and (b, c) structure of the IR_arrowhead_ in two different views. (c) Surface representation. (d) Cryo-EM map and (e, f) structure of the IR_pseudo-arrowhead_ in two orientations, shown in the same views as in (a–c) by aligning the stalk of protomer A to that of IR_arrowhead_. (g) Cryo-EM map and (h, i) structure of IR_pseudo-gamma_ in the same views as those of a–c.

The structure of the IR_arrowhead_ bound to A62D-8T closely resembles that of the A62M-bound IR (PDB 7YQ6) (24). In this conformation, the two protomers of the Λ-shaped apo IR are rotated clockwise relative to each other, pivoting at Cys524, which forms one of the disulfide linkages between the protomers (Fig. 3a–c, Supplementary Fig. 6a, b). In this symmetrical structure, the interactions between FnIII-2 and L2’ (or FnIII-2’ and L2), referred to as auto-inhibition sites, are disrupted (17). Within the binding site of the A62D-8T aptamer, we observed density corresponding to the size of only one A62M molecule in each protomer (Supplementary Fig. 6c). The reason why only one A62M binds to each IR protomer is unclear, but we speculate that the two A62M aptamers in A62D are too bulky to fit simultaneously between the L1 domain of one protomer and the FnIII-1’ domain of the other protomer. Each A62M molecule binds to opposite sides of the IR dimer, positioning itself between the L1 domain of one protomer and the FnIII-1’ domain of the other (Fig. 3b). The head of the IR_arrowhead_ is composed of the L1, CR, a portion of L2, and A62M, while the FnIII-1∼3 domains are located perpendicularly in the center, forming the stalks of the IR_arrowhead_. The distance between the membrane-proximal ends of FnIII-3 and FnIII-3’ is ∼42 Å (Fig. 3c).

### The IR_pseudo-arrowhead_ structure

The second class of particles forms a modified arrowhead-like shape, which we designate as the IR_pseudo-arrowhead_. In this asymmetrical structure, the head of one protomer (protomer A) is lifted from the symmetric IR_arrowhead_ conformation, while the opposing protomer (protomer B) remains unchanged (Fig. 3d–f). This structure was resolved at a 6.26 Å resolution (Supplementary Fig. 4).

To analyze this structure, we performed rigid-body fitting of the IR and A62M from the A62M-bound IR structure (PDB 7YQ6), which aligned well with the density. In the density map for A62D-8T, only one A62M module was visible between L1 and FnIII-1’ (or, alternatively, between L1’ and FnIII-1; Supplementary Fig. 6d). In the IR_pseudo-arrowhead_, two A62D-8T (A62M) aptamers bind to different sites on the IR dimer, which we refer to as site-u (upper side) and site-l (lower side), respectively (Fig. 3d–f and 4a, b). At site-u, the L1 domain of protomer A and the FnIII-1’ domain of protomer B simultaneously interact with the A62M aptamer. At site-l, a similar interaction is observed between the opposite protomer and a second A62M aptamer (Supplementary Fig. 6e–g). However, we hypothesize that subtle differences exist between the interactions at site-u and site-l, particularly between the residues and aptamers. In the IR_arrowhead_ conformation, both the L1 and FnIII-1 domains are involved in aptamer binding. It is possible that the A62M in the IR_pseudo-arrowhead_ conformation is slightly rotated or reoriented, leading to slight variations in the binding interface.

We superimposed the two protomers of the IR_pseudo-arrowhead_ by aligning their stalk regions (FnIII-1 to FnIII-3) (Fig. 4c). The head groups of the protomers were arranged differently, with a 13° difference when measured at Q177-L459-V604 (L1-L2-FnIII2), as protomer A is raised relative to protomer B. The distance between the E204 residues (in the CR domain) of the two aligned protomers is 32.7 Å, while the distance between their Q177 residues (in the L1 domain) is 30 Å.

**Figure 4.**
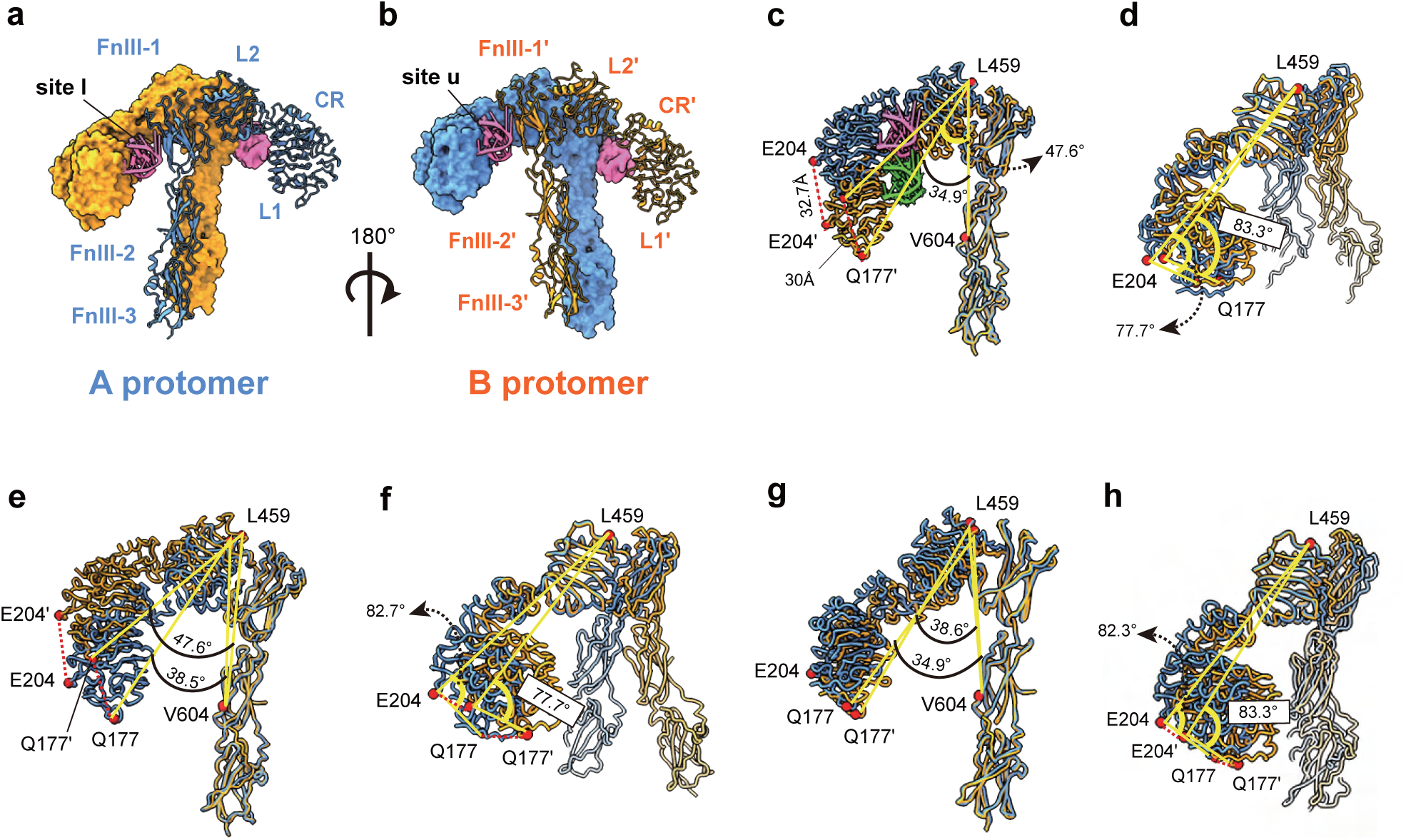
Structure of the IR_pseudo-arrowhead_ and comparison with other IR structures. (a, b) Overall structure of the IR_pseudo-arrowhead_ in two views. (c) Comparison of protomers A and B by aligning the stalks. (d) Comparison of the two protomers by aligning the L2 domain. (e) Comparison between the A protomers of IR_arrowhead_ and IR_pseudo-arrowhead_, aligned by stalks. (f) Alignment of the A protomers by the L2 domain. (g) Comparison between the B protomers of IR_arrowhead_ and IR_pseudo-arrowhead_, aligned by stalks. (h) Alignment of the B protomers by the L2 domains. IR_pseudo-arrowhead_ and IR_arrowhead_ are shown in orange and blue, respectively.

To investigate the hinge flexibility between the CR and L2 domains, we aligned the L2 domains of both protomers (Fig. 4d). In protomer A, the L1-CR domain shifts towards the L2 domain by 6°, as measured at Q177-E204-L459 (L1-CR-L2), whereas no significant changes were observed in protomer B. Consequently, the head domain of protomer A becomes more compact, with the distances between the Q177 residues and E204 residues reduced to 16 Å and 9.9 Å, respectively, in the L2-aligned protomers. In the aligned structures, the aptamers occupy slightly different positions (Fig. 4c). However, due to the shift in the L1-CR domain, the aptamer position also shifts upwards while maintaining a similar interface to that between the A62M and L1 domains in the IR_arrowhead_ structure.

### Comparison of IR_arrowhead_ and IR_pseudo-arrowhead_ structures

We compared the structures of site-u and site-l in IR_pseudo-arrowhead_ with the corresponding regions in A62M-bound IR_arrowhead_ (PDB ID: 7YQ6) by aligning their FnIII domains (Fig. 4e–h). Three major conformational differences were identified in the head arrangement of IR_pseudo-arrowhead_ relative to IR_arrowhead_. Specifically, the head domains of protomer A and B were shifted upward by 9° and downward by 4°, respectively, as measured at the Q177-L459-V604 segment (L1-L2-FnIII2; Fig. 4e, g).

In the alignment of L2, the Q177-E204-L459 (L1-CR-L2) angle differences between protomer A and protomer B were 5° and 1°, respectively, suggesting that movement at the L1-CR hinge (between CR and L2) is not substantial (Fig. 4f, h). However, the downshift of protomer A, indicated by the L1-L1’ distance (21.5 Å between Q177 residues and 18.5 Å between E204 residues), and the upshift of protomer B (9.7 Å for L1 and 11.5 Å for CR at E204) reveal that protomer A undergoes more significant changes from the IR_pseudo-arrowhead_ to IR_arrowhead_ conformation.

Additionally, both protomers display rotation relative to the IR_arrowhead_ protomer (Supplementary Fig. 6h, i). In the IR dimer, the two protomers are connected by a Cys524-Cys524’ disulfide bond, which serves as the center of rotation. Upon aligning the stalk of protomer A between IR_pseudo-arrowhead_ and IR_arrowhead_, we observed that protomer B rotates clockwise by ∼11.2° (Supplementary Fig. 6h). Due to these structural changes in the L2-FnIII1 and CR-L2 hinges, along with the rotation of protomers A and B, the distance between the membrane-proximal ends at D907-D907’ is reduced from 43.3 Å to 30.1 Å (Fig. 3b, d, Supplementary Fig. 6i).

### The IR_pseudo-gamma_ structure

The third class of particles displayed more pronounced conformational differences from the IR_arrowhead_ structure compared to the IR_pseudo-arrowhead_ structure (Fig. 3g–i, 5a, b, and Supplementary Fig. 5). In this conformation, the L1 domain of protomer A is shifted upward, positioned almost perpendicular to the stalk, while the L1’ domain of protomer B is shifted downward, closely aligned with the stalk. This third-class conformation resembles the fully active Γ-shape observed in the insulin-bound or insulin and A43 aptamer-bound forms of the receptor (Fig. 5b, c).

**Figure 5.**
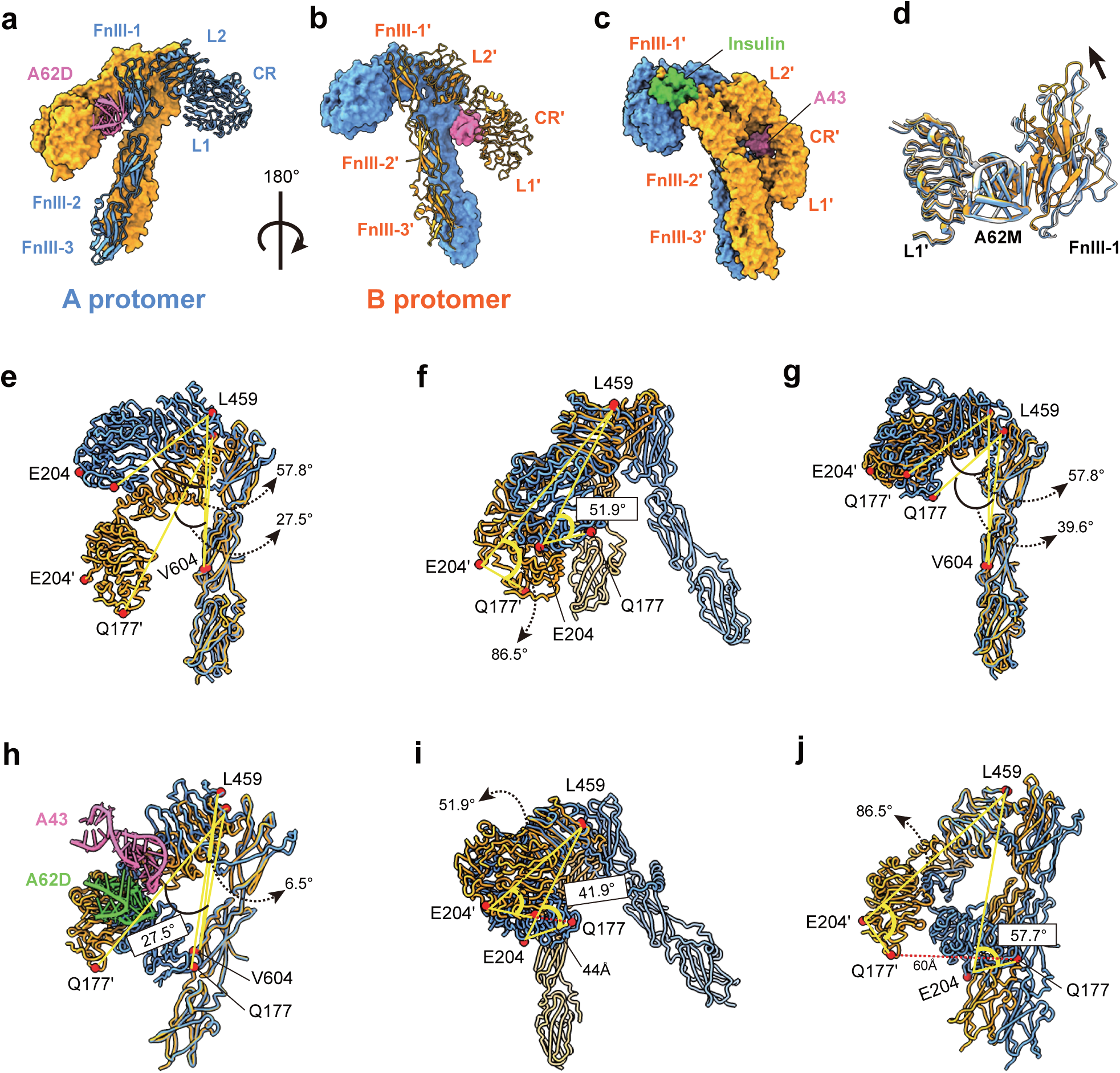
Structure of IR_pseudo-gamma_ and comparison with other IR structures. (a, b) Two views of IR_pseudo-gamma_. Protomers A and B are shown in blue and orange, respectively. (c) Structure of IR_gamma_ in the same orientation as in (b). (d) Superimposed structures of the A protomers of IR_pseudo-gamma_ (orange), IR_pseudo-arrow_ (blue) and IR_arrowhead_ (gray), aligned by the A62M, showing the relative positions of the FnIII-1 domains between the two protomers. (e) Comparison of protomer A and B of IR_pseudo-gamma_ by aligning the stalks. IR_pseudo-gamma_ and IR_gamma_ are shown in orange and blue, respectively. (f) Comparison of protomer A and B of IR_pseudo-gamma_ by aligning the L2 domain. (g, h) Superimposed structures of the A (g) and B (h) protomers of IR_pseudo-gamma_ and IR_gamma_, aligned by the FnIII stalks. (i, j) Superimposed structures of the A (i) and B (j) protomers of IR_pseudo-gamma_ and IR_gamma_, aligned by the L2 domains.

Unlike the IR_arrowhead_ or IR_pseudo-arrowhead_ conformations, A62D binds at only one site, specifically at the interface between the L1 domain of protomer B and the FnIII-1’ domain of protomer A (Fig. 3g–i). The interaction between the L1 domain and the aptamer is similar to the binding observed between IR_arrowhead_ and A62M (Fig. 3b, 5b). However, the orientation of the FnIII-1 domain in the IR_pseudo-gamma_ conformation differs from that in the IR_pseudo-arrowhead_ and IR_arrowhead_ structures (Fig. 5d). Comparison of the structures between IR_pseudo-gamma_ and IR_arrowhead_ or IR_pseudo-arrowhead_ is described in supplementary information (Supplementary Fig 7).

A structural comparison of protomers A and B, aligned by their stalk regions, reveals that the head of protomer A is tilted upward by 30° compared to that of protomer B, as measured by the angles formed by Q177-L459-V604 (L1-L2-FnIII2); 58° versus 27°, respectively (Fig. 5e). When the L2 domains are aligned, protomer A exhibits a more compact head domain, with an angle of 52° (Q177-LE204-L459), while the head of protomer B adopts a more extended conformation, with a corresponding angle of 86.5° (Fig. 5f). The relative distances between the Q177 (L1) residues and the E204 (CR) residues of the two protomers are 60.1 Å and 41 Å, respectively. Thus, aptamer binding induces a more extended conformation of the head domain in protomer B.

### Structural comparison of IR_pseudo-gamma_ and IR_gamma_

The fully activated conformation of IR_gamma_ is achieved when bound by a single insulin molecule, or by insulin and A43 (24, 47), with one side of the receptor’s head (protomer A) lifted to accommodate insulin between the L1, CR, and FnIII-1’ domains, while the opposite side (protomer B) remains insulin-free and shifts downward to form a compact head structure (Fig. 5c). Although it is uncertain whether IR_pseudo-gamma_ represents an active conformation, the proximity of the membrane-proximal ends of the FnIII-stalks (∼20 Å apart) suggests it might correspond to a fully phosphorylated state (Fig. 3i).

When aligning the stalk of protomer A in IR_pseudo-gamma_ with that of IR_gamma_, we observed a downward shift in the aptamer-free head of IR_pseudo-gamma_ and an upward shift in the aptamer-bound head (or vice versa) compared to IR_gamma_ (Fig. 5g, h). This movement is further demonstrated by changes in specific domain angles. The Q177-L459-V604 (L1-L2-FnIII2) angle decreased from 58° in IR_pseudo-gamma_ to 40° in IR_gamma_ (Fig. 5g). Additionally, the L1 domain in protomer A of IR_pseudo-gamma_ was displaced by 26.4 Å, measured at residue Q177, compared to its position in IR_gamma_.

In protomer B of IR_pseudo-gamma_, the Q177-L459-V604 angle was 27.5°, much larger than the 6.5° observed in IR_gamma_ (Fig. 5h). Similarly, the E204-L459-V604 angle in IR_pseudo-gamma_ was 39.1°, significantly greater than the 13.2° in IR_gamma_. These differences indicate that the head domain of protomer B in IR_pseudo-gamma_ adopts a more extended conformation compared to that of IR_gamma_.

When superimposing the L2 domains, the Q177-E204-L459 angle in protomer A of IR_pseudo-gamma_ was 52°, slightly wider than the 42° angle seen in IR_gamma_ (Fig. 5i). For protomer B, the head domain in IR_pseudo-gamma_ was notably more open and extended, with an angle of 86.5° compared to 57.7° in IR_gamma_ (Fig. 5j). The distance between L1 domains at Q177 was measured at 44 Å for protomer A and 60 Å for protomer B when the L2 domains were aligned. These findings indicate significant conformational changes in both the CR-L2 and L2-FnIII1 hinges.

### A62D-induced crosslinking of IR dimers

Our structural analysis demonstrated that A62D induces the formation of three distinct IR conformations, one of which closely resembles the fully active IR conformation. This structure likely contributes to the full phosphorylation of IR. Given that the IR_pseudo-gamma_ dimer binds only a single A62M module of A62D, we hypothesize that each A62M unit binds separately to an IR dimer, facilitating cross-interaction between the dimers. This interaction positions the dimers in close proximity, promoting trans-phosphorylation. This hypothesis is supported by the data presented in Fig. 1b, which shows that A62D linked with a connector of 8T to 19T efficiently phosphorylated IR. In contrast, shorter or longer linkers failed to achieve full phosphorylation. This may be due to short linkers bringing the IR dimers too close together, limiting the flexibility required for trans-phosphorylation, while longer linkers keep the dimers too far apart to efficiently facilitate intermolecular trans-phosphorylation. Therefore, we conclude that A62D aptamers with optimally sized linkers can effectively position two IRs in close proximity, inducing phosphorylation of the IR dimers.

To further validate this model, we mixed A62M or A62D with IR dimers in 4:1 and 2:1 molar ratios, respectively, and conducted size-exclusion chromatography analysis (Fig. 6a, b). We observed substantial fractions of oligomeric A62D-IR complexes in the void volume, whereas relatively small fractions of A62M-IR appeared in this region. Similar amounts of 1:1 A62M-IR complexes were observed. Additionally, negative staining analysis revealed numerous oligomeric A62D-IR particles in the void volume fractions (Fig. 6c–f).

**Figure 6.**
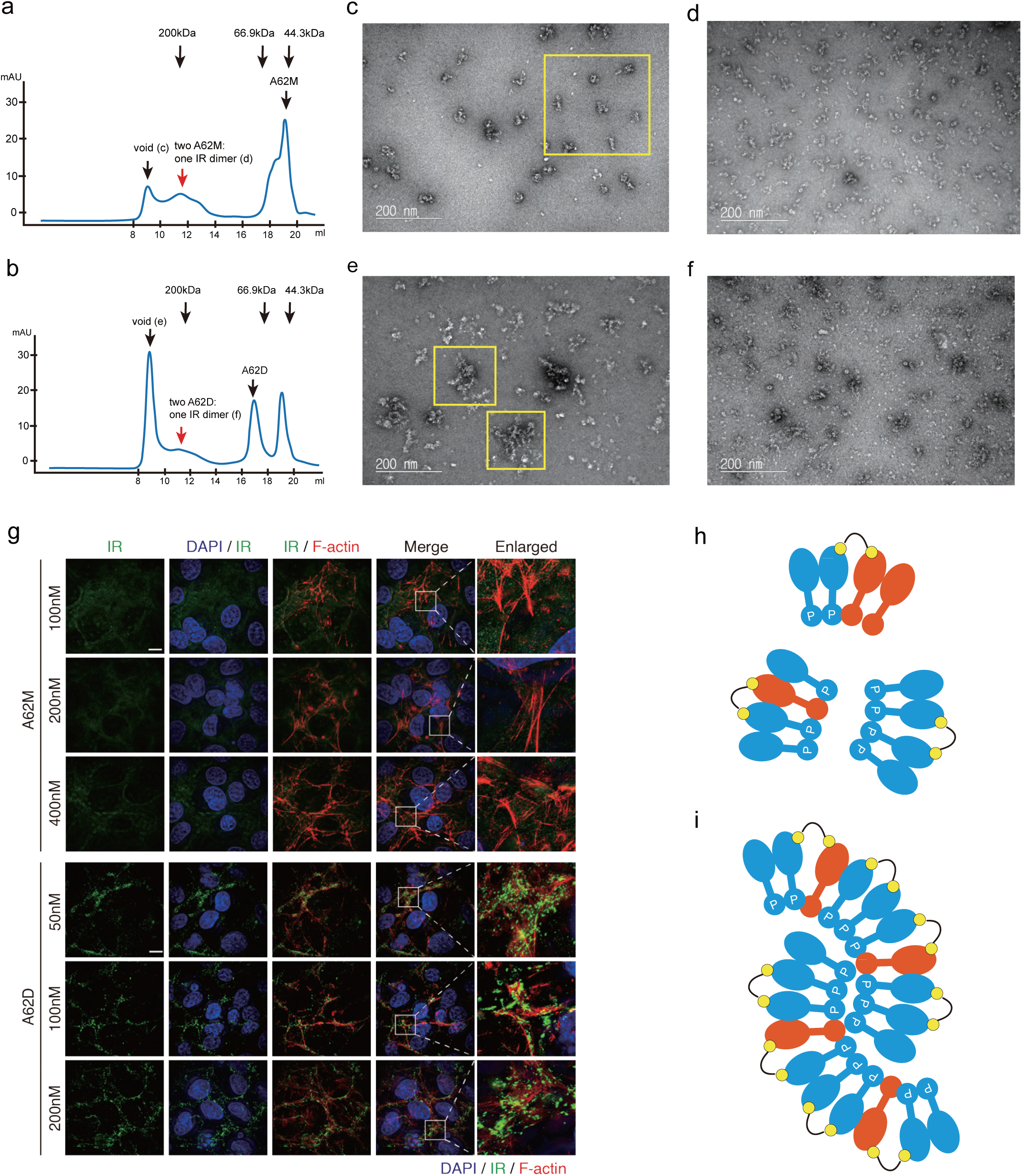
Intermolecular interactions of A62D-induced IR dimers. (a, b) Comparison of the profiles of IR mixed with A62M and A62D aptamers. (c-f) Comparison of negative staining images of fractions from the void volume peak (c) and normal peak (d) of the A62M-IR complex, and fractions from the void volume peak (e) and normal peak (f) of the A62D-IR complex. (g) Representative images from structured illumination microscopy (SIM). Rat-1/hIR cells treated with indicated aptamers at various concentrations were co-immunostained with anti-insulin receptor (IR), DAPI and phalloidin (F-actin). Scale bar, 10mm. (h) A model of the A62D complex with two IR dimers. (i) A model of the “beads on a string”-like cluster formed by A62D complexes with IR dimers.

We also examined whether A62D induced the oligomerization of IR in live cells. We stained the IR using a fluorescent-labeled antibody, and observed recruitment of IR by A62D in live cells using super-resolution fluorescence microscopy (structured illumination microscopy). Rat-1/hIR cells expressing IR were treated with A62M and A62D at various concentrations (100nM, 200nM, 400nM for A62M; 50nM, 100nM, 200nM for A62D) and IR oligomerization was monitored (Figure 6g). While we did not observe noticeable IR clusters in A62M-treated cells, significant number of IR clusters were observed in A62D-treated cells (Figure 6g, Supplementary Fig 8a). Although IR oligomerization in A62D-treated cells was more prominent in the plasma membrane, marked by F-actin, clustering events were also elevated in the cytoplasm and nucleus (Figure 6g, Supplementary Fig 8b–d). Collectively, these results demonstrate that A62D aptamer induces full IR phosphorylation by facilitating intermolecular interactions between the IR dimers (Fig. 6h–i).

## Discussion

In this study, we aimed to enhance the previously reported A62M aptamer, a partial agonist known to induce Tyr1150 mono-phosphorylation of IR (24, 31). Our findings revealed that crosslinking two A62M aptamers with linkers of varying lengths resulted in full phosphorylation of the IR and increased potency. To our knowledge, this represents an unprecedented example of aptamer-mediated full activation of the IR.

The primary question raised by our study is why the A62D-8T (and other A62D variants with different linkers) behaves differently from A62M, leading to full phosphorylation of the IR. We hypothesized that A62D induces a distinct structural change in the IR compared to A62M. To explore this, we determined the structure of the A62D-bound IR and compared it with the A62M-IR complex. We observed three conformations induced by A62D: the previously identified arrowhead conformation, along with pseudo-arrowhead and pseudo-gamma conformations. In all three structures, only one A62M module of the A62D binds to the IR (Fig. 3b, e, h). The A62M modules bind at the interface between the L1-CR region of one IR protomer and the FnIII-1’ region of another protomer. It appears that the full A62D structure is too bulky to fit between these regions.

While the IR_pseudo-arrowhead_ structure closely resembles the IR_arrowhead_ conformation, the IR_pseudo-gamma_ conformation is distinct from IR_arrowhead_ and more closely resembles the IR_gamma_ structure observed in the presence of a single insulin molecule or a single insulin molecule with the A43 positive allosteric modulator (24, 47). Although the activation state of IR_pseudo-gamma_ remains unclear, its structural similarity to IR_gamma_ and the proximity of the membrane-proximal ends of the FnIII stalks suggest that IR_pseudo-gamma_ could contribute to full activation of the IR. Initially, based on structural analysis, we considered a possibility that IR_pseudo-gamma_ plays a key role in the A62D-induced full phosphorylation of the IR. The particle distribution for IR_arrowhead_, IR_pseudo-arrowhead_, and IR_pseudo-gamma_ in the A62D complex was observed at a ratio of 0.29:0.3:0.12, respectively. Despite the relatively small proportion of IR_pseudo-gamma_, assuming that this conformation is fully phosphorylated, we estimated that the presence of 12% fully phosphorylated IR in the presence of A62D, compared to none in the A62M-bound state, could be sufficient for detection in our system (Fig. 1c). To determine whether IR_pseudo-gamma_ is exclusive to the A62D-bound state, we reanalyzed the A62M-bound IR data and found that ∼7% of the particles also exhibited the IR_pseudo-gamma_ conformation. Thus, while we cannot rule out the possibility that IR_pseudo-gamma_ contributes to full phosphorylation, it is unlikely to be the primary mechanism by which A62D induces full IR phosphorylation.

We then considered an alternative explanation: A62D, containing two A62M modules, may bring two or more IR dimers into close proximity, facilitating intermolecular trans-phosphorylation. This hypothesis is supported by increased oligomerization observed in the presence of A62D compared to A62M, as shown by size exclusion chromatography and negative staining. The length of the linker between the two A62M modules is crucial in this crosslinking process. If the linker is too short, each A62M module may not efficiently capture the IR dimers; if it is too long, the bound IR dimers may not interact efficiently enough for optimal intermolecular phosphorylation.

We note that our single-particle cryo-EM analysis revealed the structures of the A62D-bound IR dimer because we focused on the fraction containing homogeneous particles representing a single IR dimer. Although the structural analysis identified three distinct conformations of the A62D-bound single IR dimer, biochemical and negative staining analyses suggest that intermolecular interactions between IR dimers are essential for full phosphorylation and activation of the IR.

A62D may promote the oligomerization of IR dimers through two potential mechanisms. First, each A62D could recruit two IR dimers, one at each end. In this scenario, only one protomer of an IR dimer interacts with the A62M module of A62D, likely positioning the IR in the IR_pseudo-gamma_ conformation (Fig. 6h). Alternatively, multiple IR dimers could be continuously linked via several A62D aptamers in a “beads-on-a-string” configuration (Fig. 6i). In this case, each A62D would bind to IR_pseudo-arrowhead_ or IR_arrowhead_ conformations at both ends. The latter model predicts a higher proportion of oligomerized IR dimers in the void volumes during size-exclusion chromatography (Fig. 6b, e, f, i). However, it is possible that both models coexist in A62D-induced IR oligomers.

Conventional models have depicted IR activation as a simple switch dependent on insulin binding. However, recent studies highlight the importance of spatial dynamics of IR at the plasma membrane in receptor activation and signal transduction. Super-resolution microscopy has shown that IR molecules are incorporated into dynamic clusters (48, 49). In this model, IR dimers are recruited into clusters at the plasma membrane, cytoplasm, and nucleus, with insulin stimulation further promoting cluster formation in their active states. This results in an increase in both the number of IR-containing clusters and the number of IR molecules per cluster. Additionally, rod-like insulin–DNA origami nanostructures have demonstrated that increasing insulin valency enhances both IR and AKT phosphorylation (50). These findings suggest that the formation of IR dimers or oligomers plays a critical role in insulin-induced IR activation and downstream signaling, supporting our interpretation of A62D-induced IR activation. However, the precise mechanisms by which IR clustering enhances receptor phosphorylation and signaling remain unclear.

Over the past few decades, various agonists that activate the insulin receptor have been developed using antibodies, peptides and aptamers (10, 12, 17, 20–22, 28, 31, 32, 51–55). These agonists exhibited high potency in activating IR. However, because their potencies have been studied through different approaches with A62D, direct comparisons of their potency on IR with that of A62D are not straightforward. Our study indicates that A62D-8T exhibits a slow and sustained activation of IR, while it induces up to a two-fold increase in IR and AKT phosphorylation compared to insulin. A common feature of several agonists in a manner distinct from insulin is that they selectively activate the metabolic functions and ATK pathway of the insulin receptor. Such functional selectivity appears to be associated with site specific phosphorylation and specific structural state of IR. In particular, S597, an insulin mimetic peptide, and A62M induce selective phosphorylation at Y1150 of the insulin receptor and nearly identical arrowhead structures (24, 53). Based on these studies, the stepwise activation model has been proposed, suggesting that the function of the insulin receptor is selectively regulated according to its conformational states (14).

Despite these advances, previous studies on the agonists have described that a single insulin receptor acts in a stand-alone state, and have not explained the functional role of insulin receptor clustering on the cell membrane. In this study, we artificially induced clustering of IR using A62D, and demonstrated that insulin receptor activation can be regulated by inter-receptor interactions. Collectively, these studies suggest that spatial clustering strengthens intermolecular interactions between IR molecules, leading to full receptor phosphorylation and the selective enhancement of AKT pathway. Furthermore, A62D suggests a new possibility for diabetes treatment. Investigating the potential clinical effects of artificially induced IR clustering in diabetes patients is an intriguing topic for future research. However, optimizing the aptamer to effectively induce IR clustering under in vivo conditions while maintaining sufficient pharmacokinetics and pharmacodynamics remains a major challenge.

While the structural transition in IR activation involves a shift from the inactive, Λ-shaped dimer to the active, single-insulin-bound IR_gamma_ conformation, the structures of the intermediate states between these two conformations remain poorly understood. Recent studies have proposed potential intermediates, such as the tilted T-shaped IR, resulting from binding two or more insulin molecules or other ligands (9, 10, 22–24). However, these structures require further investigation. All conformations observed in this study arise from hinge-bending motions between the L2 and FnIII-1 domains and between the CR and L2 domains, along with rigid-body rotations of the two protomers in the Λ- and gamma-shaped IR dimer. Therefore, the A62D-induced IR structures presented here may offer insights into the intermediate states involved in IR activation.

In summary, by crosslinking A62M, we enhanced the efficacy of the IR aptamer, achieving full activation of the receptor. This study presents a potential strategy for designing more effective IR agonists as therapeutics. By increasing intermolecular interactions between IR dimers, agonist-induced IR signaling could be more efficiently transduced.

## Acknowledgments

We thank Photon Science Center at PAL for the help of data collection. This work was supported by grants from the National Research Foundation of Korea (NRF) funded by the Korea government (MEST, No. 2021R1A2C301335711, RS-2024-00344154, and MSIT, Bio&Medical Technology Development program RS-2024-00440289 to Y.C., No. 2021R1A2C301335711 and RS-2024-00344154 to J.K., NRF-2022R1A2C1091474 to E.J.O.), the Korea-US Collaborative Research Fund (KUCRF), funded by the Ministry of Science and ICT and Ministry of Health & Welfare, RS-2024-00466776 to H.L., NRF funded by the Ministry of Education (RS-2023-00241226) and new professor research program of KOREATECH to N, Y., Basic Science Research Institute Fund (2021R1A6A1A10042944) to J. K. and the BK21 program (Ministry of Education) to Y.C., J.K, and H.N. The Structured Illumination Microscopy images were acquired using the AP DeltaVision OMX Ultra High-Resolution Fluorescence Microscope at the SNU Center for Macromolecular and Cell Imaging (SNU-CMCI).

## Author contributions

J.K. carried out protein expression, purification, and structure determination with the help of H.N.; N.Y. and H.N. performed biochemical experiment with the help of E.J.O.; S-Y.C. and H.L. performed in vivo imaging analysis; J.K., N.Y., S.H.R. and Y.C. designed the research; N.Y. and Y.C. wrote the manuscript with the help of J.K., H.N., S.H.R.

## Data availability

Atomic coordinates and the cryo-EM map have been deposited in the PDB and the EM Data Bank, respectively, under following accession numbers; IR_arrowhead_ (EMD-61490, PDB 9JHS), IR_pseudo-arrowhead_ (EMD-61431, PDB 9JF9); IR_pseudo-gamma_ (EMD-61432, PDB 9JFD).

## Supplementary Information

**Structural comparison of IR_pseudo-gamma_ with IR_pseudo-arrowhead_ and IR_arrowhead_**

To investigate the structural changes from IR_pseudo-arrowhead_ to IR_pseudo-gamma_, we aligned the stalks of protomer A (Supplementary Fig. 7a). In IR_pseudo-gamma_, the aptamer-free head (L1-CR domain) of protomer A is elevated by 10.2° compared to the aptamer-bound protomer A of IR_pseudo-arrowhead_ (Q177-L459-V604; L1-L2-FnIII2). Meanwhile, the aptamer-bound protomer B in IR_pseudo-gamma_ is downshifted by 7.2° relative to protomer B in IR_pseudo-arrowhead_ (Supplementary Fig. 7b). The L1 domain of protomer A in IR_pseudo-gamma_ has shifted by 29.5 Å, while in protomer B, it has shifted by 16.4 Å.

Alignment of the L2 domains between IR_pseudo-gamma_ and IR_pseudo-arrowhead_ revealed that protomer A of IR_pseudo-gamma_ adopts a more compact form (Q177-E204-L459; L1-CR-L2, 52°) compared to protomer A of IR_pseudo-arrowhead_ (78°). In contrast, protomer B of IR_pseudo-gamma_ maintains a similar conformation to protomer A of IR_pseudo-arrowhead_ (86° vs. 83°) (Supplementary Fig. 7c, d). The distance between L1 domains in protomer A and B of IR_pseudo-gamma_ is 43.3 Å and 6.7 Å, respectively. Overall, protomer A undergoes significant changes at two hinge regions (CR-L2 and L2-FnIII1), while protomer B shows relatively smaller changes, particularly in the L2-FnIII1 hinge.

When aligning the stalk of IR_pseudo-gamma_ with that of IR_arrowhead_, one head shifts upward while the other shifts downward at the L2-FnIII1 hinge (Supplementary Fig. 7e, f). The ligand-free protomer A is uplifted by 19.3° and 15° at Q177-L459-V604 (L1-L2-FnIII2) and E204-L459-V604 (CR-L2-FnIII2), respectively, while the aptamer-bound protomer B is downshifted by 1.1° and 10.6°. The L1 domain of protomer A shifts by 48 Å, while in protomer B, it shifts by 26 Å.

Upon aligning the L2 domains, the aptamer-free protomer A of IR_pseudo-gamma_ forms a more compact structure (angle reduced from 82° to 52°), while the aptamer-bound protomer B retains a nearly identical conformation (angle shift from 82° to 86°) (Supplementary Fig. 7g, h). The distance between L1-L1’ or CR-CR’ is much smaller for protomer B (5 Å) compared to protomer A (60 Å and 44 Å, respectively). Thus, protomer A experiences substantial shifts at two hinges, while the L1-FnIII1 hinge of aptamer-bound protomer B undergoes only minor adjustments.

## Supplementary Figure Legends

**Supplementary Figure 1.**
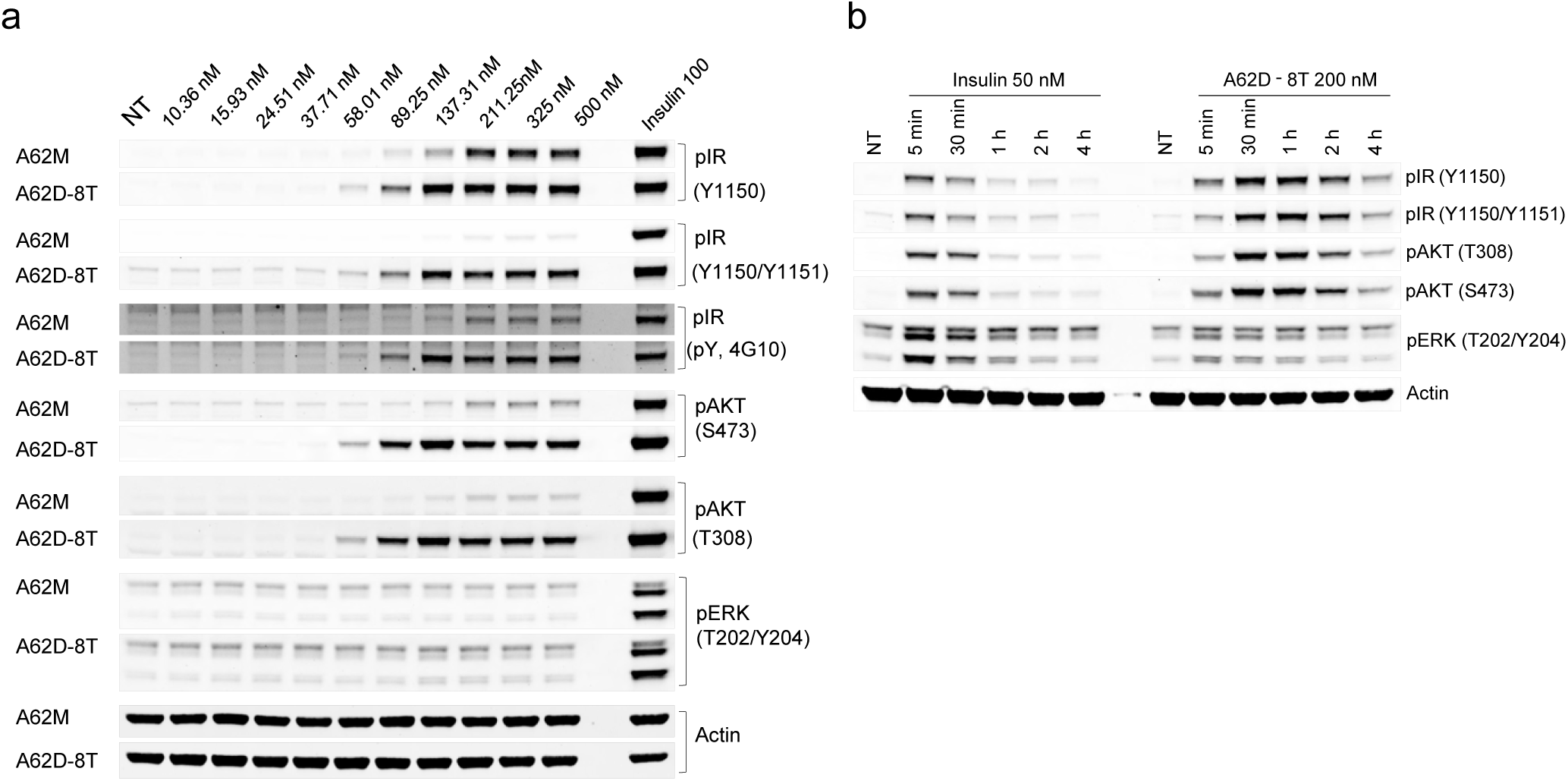
Effects of the dose and duration of A62M and A62D-8T on IR phosphorylation and downstream signaling. (a) Comparison of dose-dependent phosphorylation of IR and downstream signaling by A62M or A62D-8T. Rat-1/hIR cells were incubated with varying concentrations of A62M or A62D for 1 h. (b) The relative band intensities, compared to 100 nM insulin, are shown for pIR Y1150, pIR Y1150/Y1151, pAKT T308, pAKT S473, and pERK T202/Y204. Quantified values for the band intensities are shown in Fig 2.

**Supplementary Figure 2.**
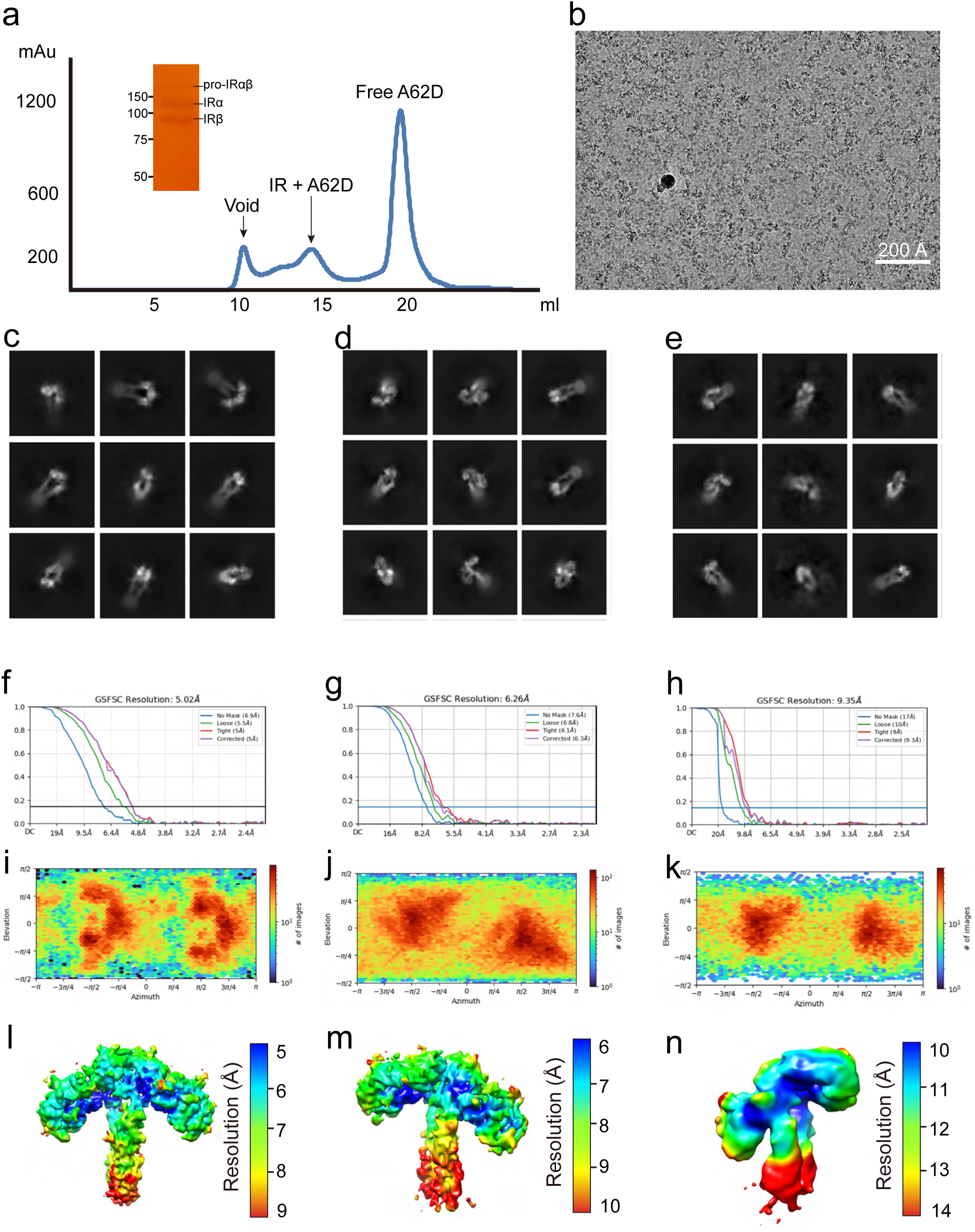
Purification of the IR + A62D complex, analysis of the quality of the cryo-EM map. (a) Size-exclusion chromatography profile and SDS-PAGE result. (b) Representative cryo-EM micrograph of the IR + A62D complex. **c-e,** 2D class averages of (c) IR_arrowhead_, (d) IR_pseudo-arrowhead_ and (e) IR_pseudo-gamma_. **f-n**, Fourier shell correlation curves (**f-h**), Angular distributions (**i-k**) and Local resolution of the map (**l-n**) of (**f, i, l**) IR_arrowhead_, (**g, j, m**) IR_pseudo-arrowhead_ and (**h, k, n**) IR_pseudo-gamma_.

**Supplementary Figure 3.**
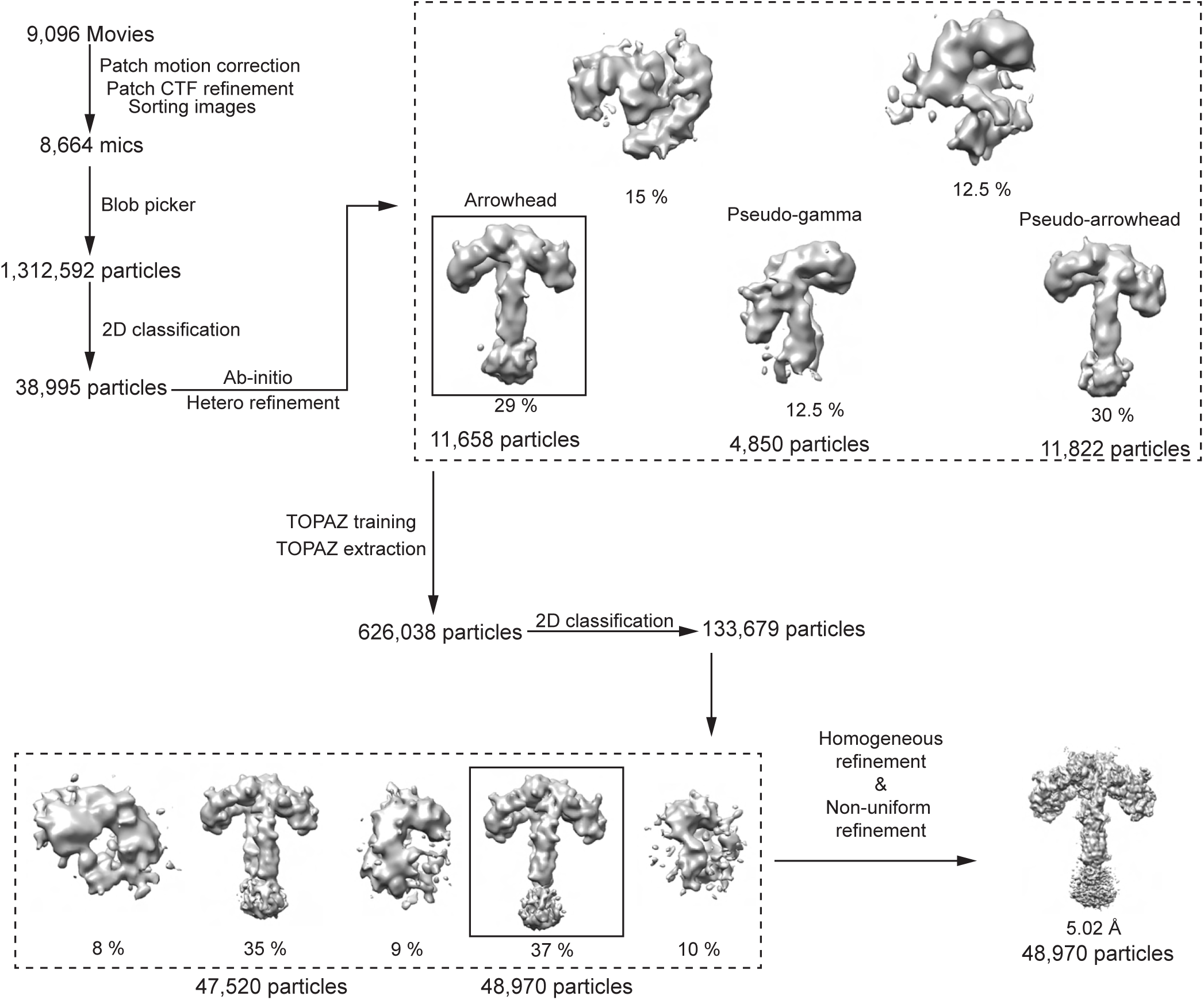
Workflow of cryo-EM processing for the IR_arrowhead_. Flow chart of data processing for the IR_arrowhead_.

**Supplementary Figure 4.**
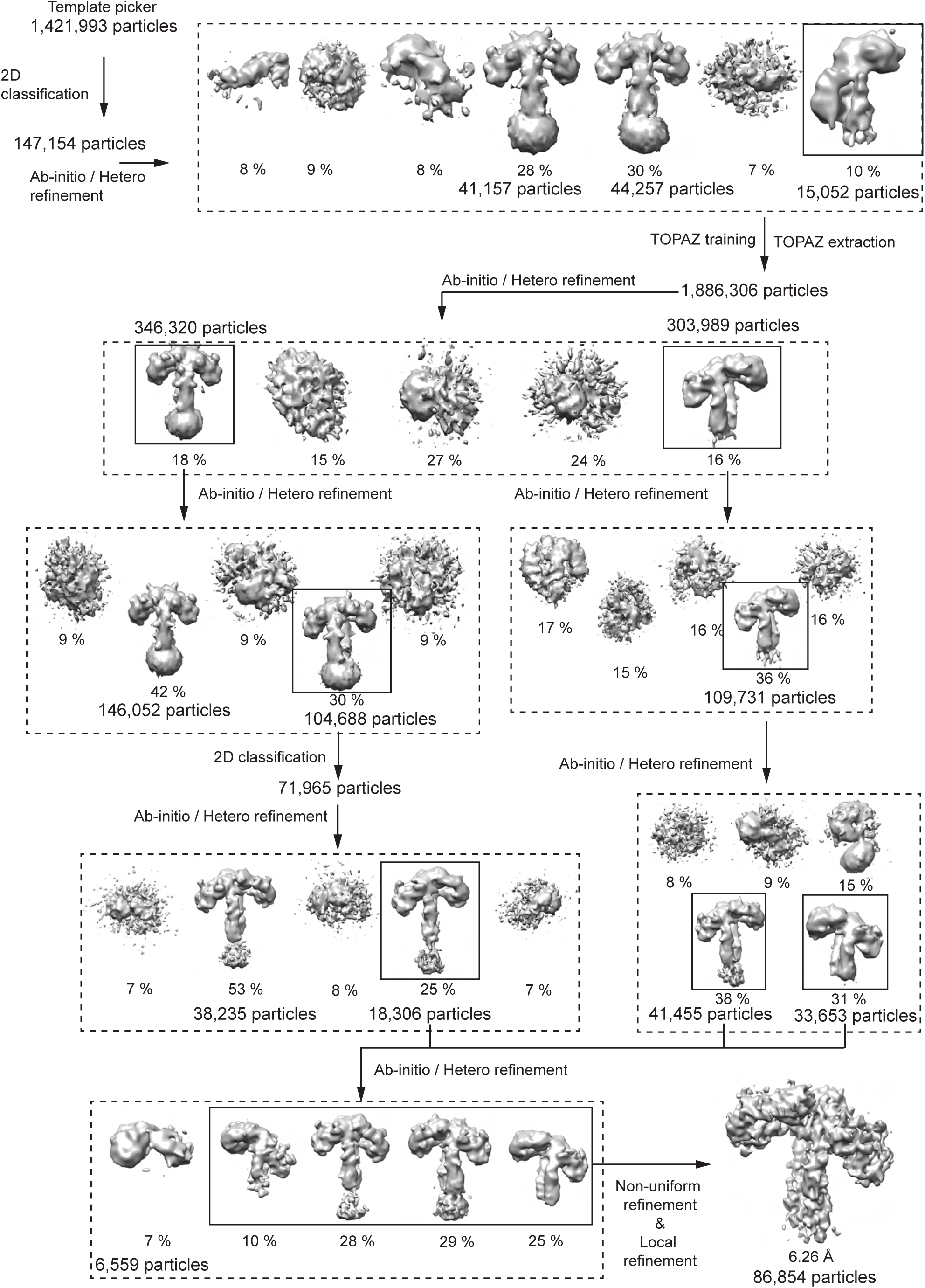
Workflow of cryo-EM processing for the IR_pseudo-arrowhead_.

**Supplementary Figure 5.**
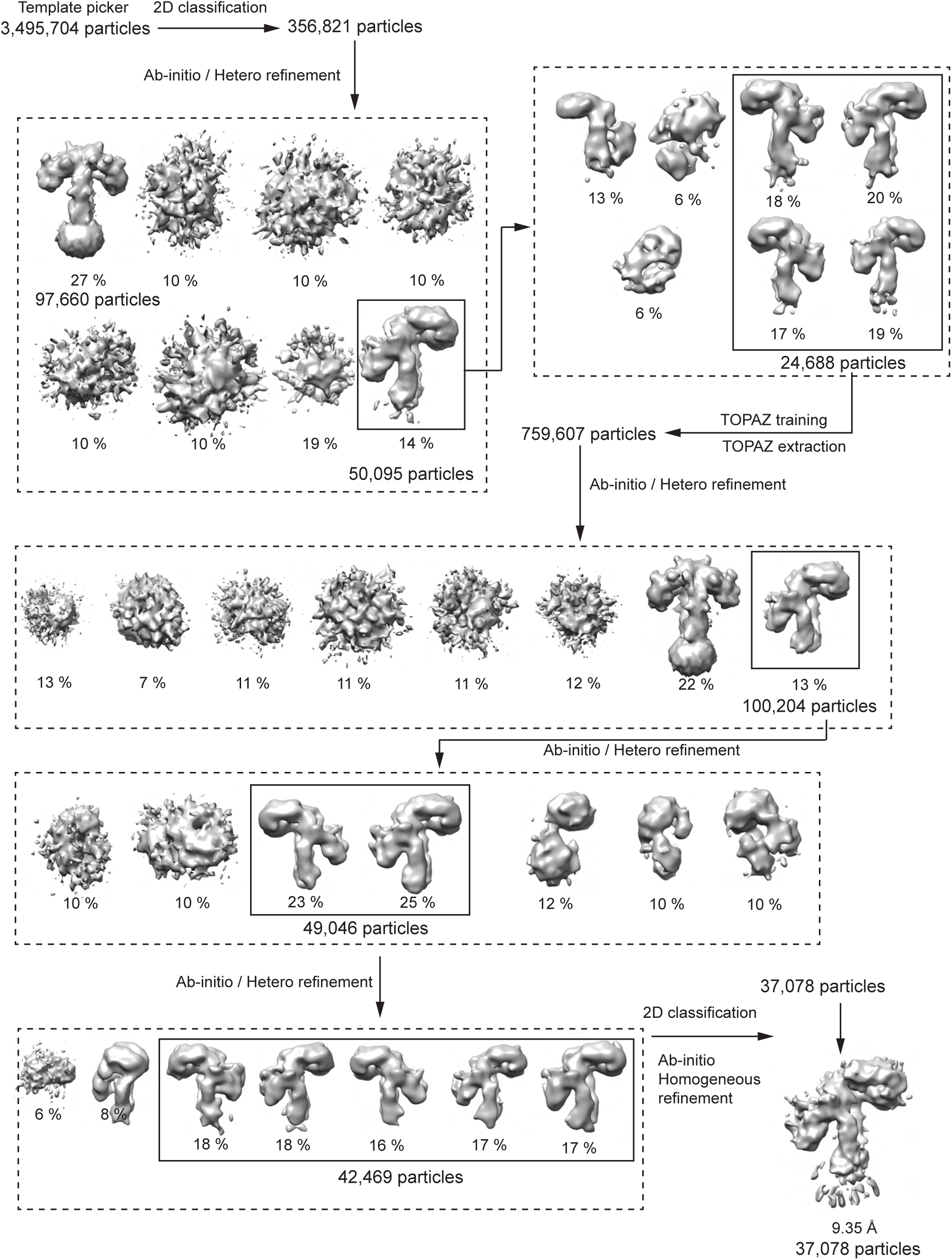
Workflow of cryo-EM processing for the IR_pseudo-gamma_.

**Supplementary Figure 6.**
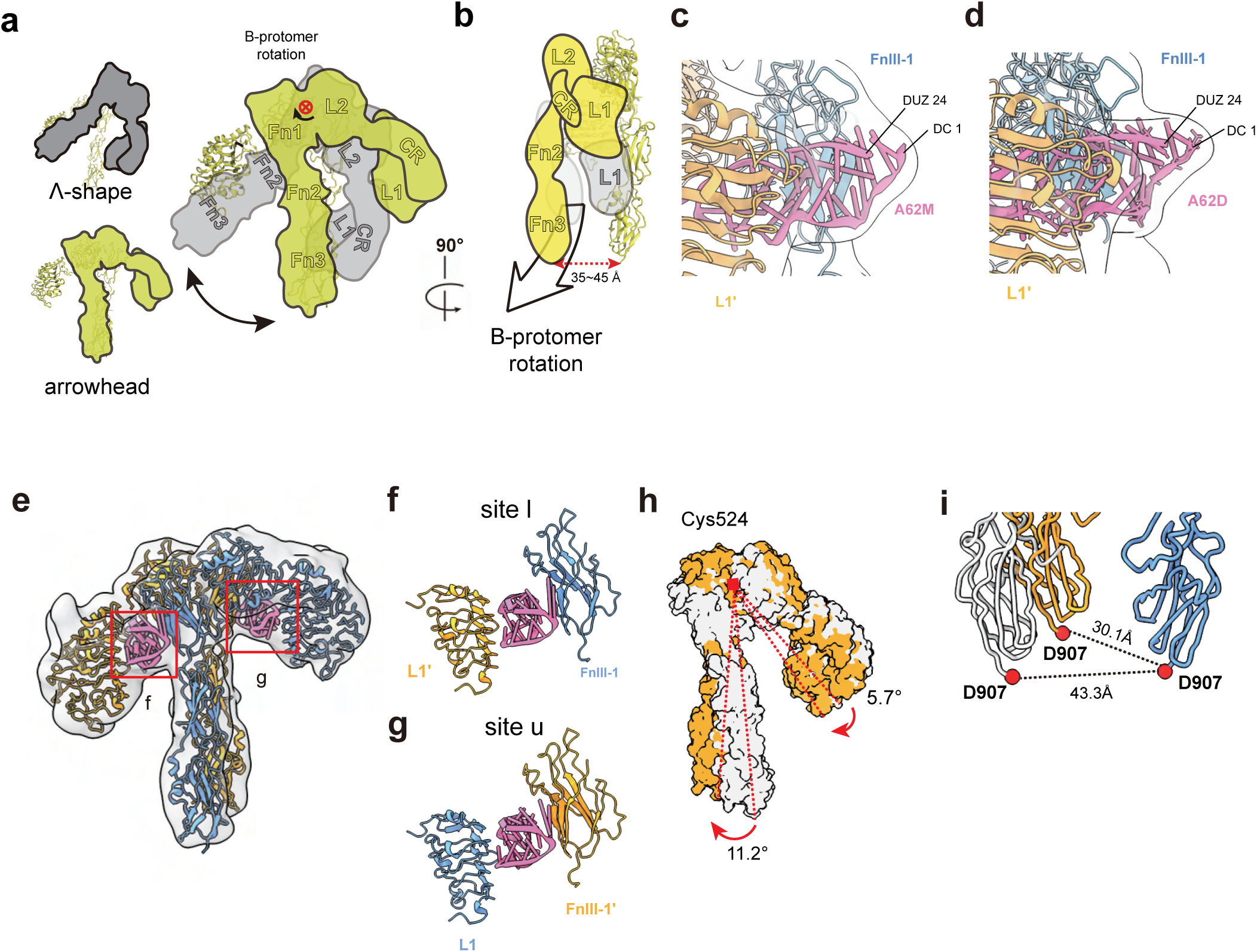
Close-up view of the aptamer binding site in IR_pseudo-arrowhead_. (a) A cartoon representation of the rigid-body rotation of the IR_apo_ (gray) to the IR_arrowhead_ (orange). Only one protomer is shown for clarity. (b) Rigid-body rotation in a is shown in a 90°-rotated view. (c) A close up view of the cryo-EM density for the A62D in the IR_pseudo-arrowhead_. Only one A62M module of the A62D can be modeled into the density. (d) Cryo-EM density for the A62D in IR_pseudo-arrowhead_. (e) Interaction of A62M with the L1 and FnIII-1 domains at site l and site u in IR_pseudo-arrowhead_. (f) Close-up view of the site-l interface, (g) Close-up view of the site-u interface. (h) Superimposed structures of the IR_arrowhead_ and IR_pseudo-arrowhead_ by aligning protomer A. Comparison of the B protomers between IR_arrowhead_ and IR_pseudo-arrowhead_ illustrates the rigid-body rotation of protomer B. (i) Aligned structures of IR_arrowhead_ (white) and IR_pseudo-arrowhead_ (yellow, blue). Distance between the membrane-proximal ends of the FnIII-3 stalks (D907-D907’) is shown for both IRs.

**Supplementary Figure 7.**
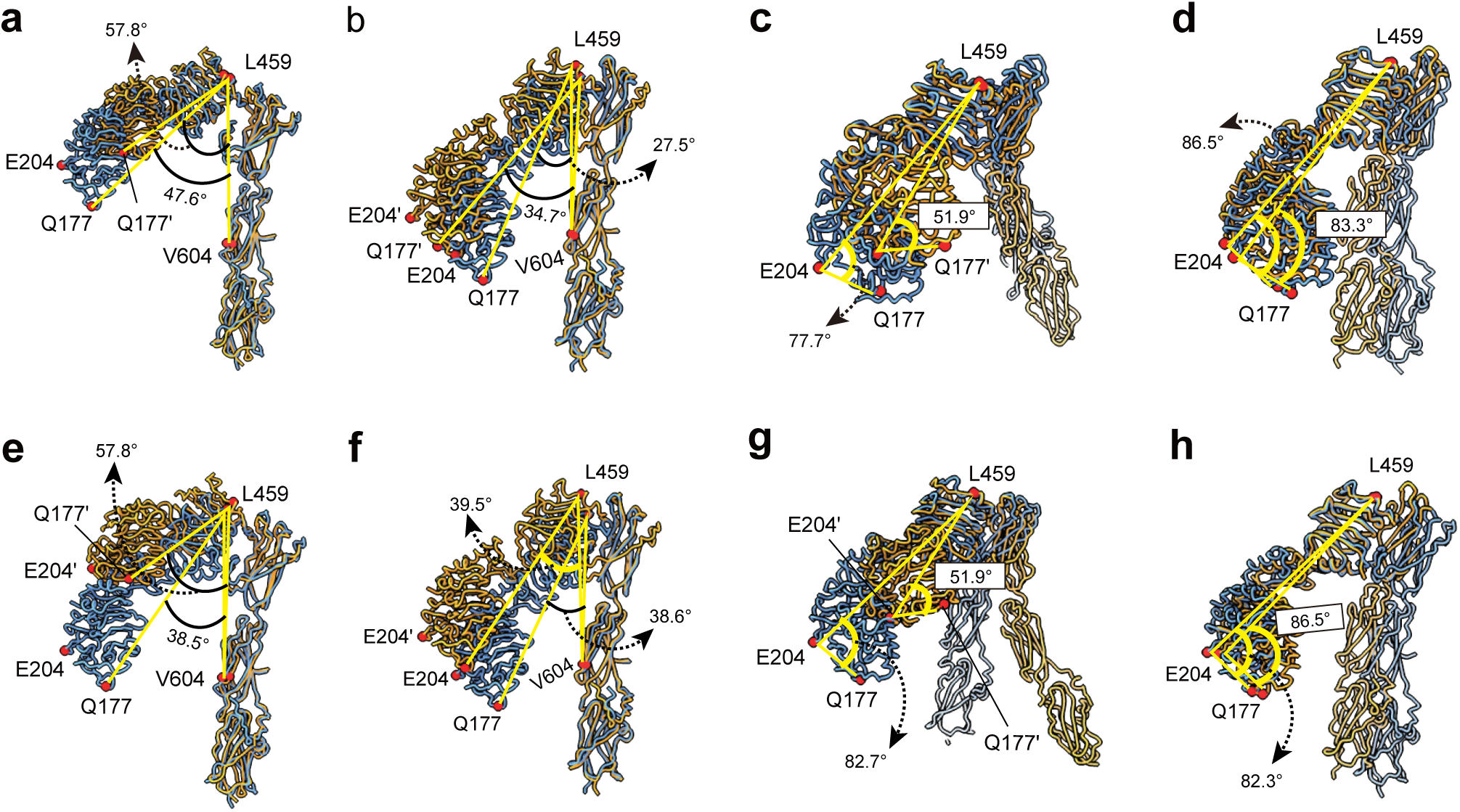
Structural comparison of IR_pseudo-gamma_ with IR_pseudo-arrowhead_ and IR_arrowhead_. (a, b) Superimposed structures of protomer A (a) and protomer B (b) in IR_pseudo-gamma_ (orange) with the respective protomers in IR_pseudo-arrowhead_ (blue), aligned by stalks. (c, d) Superimposed structures of protomer A (c) and protomer B (d) in IR_pseudo-gamma_ (orange) with the respective protomers in IR_pseudo-arrowhead_ (blue) aligned by the L2 domains. Protomers of the IR_pseudo-gamma_ and IR_pseudo-arrowhead_ are shown in orange and blue, respectively. (e, f) Superimposed structures of protomer A (e) and protomer B (f) in IR_pseudo-gamma_ (orange) with the respective protomers in IR_arrowhead_ (blue), aligned by stalks. (g, h) Superimposed structures of protomer A (g) and protomer B (h) in IR_pseudo-gamma_ with the respective protomers in IR_arrowhead_, aligned by the L2 domains. Protomers of the IR_pseudo-gamma_ and IR_arrowhead_ are shown in orange and blue, respectively.

**Supplementary Figure 8.**
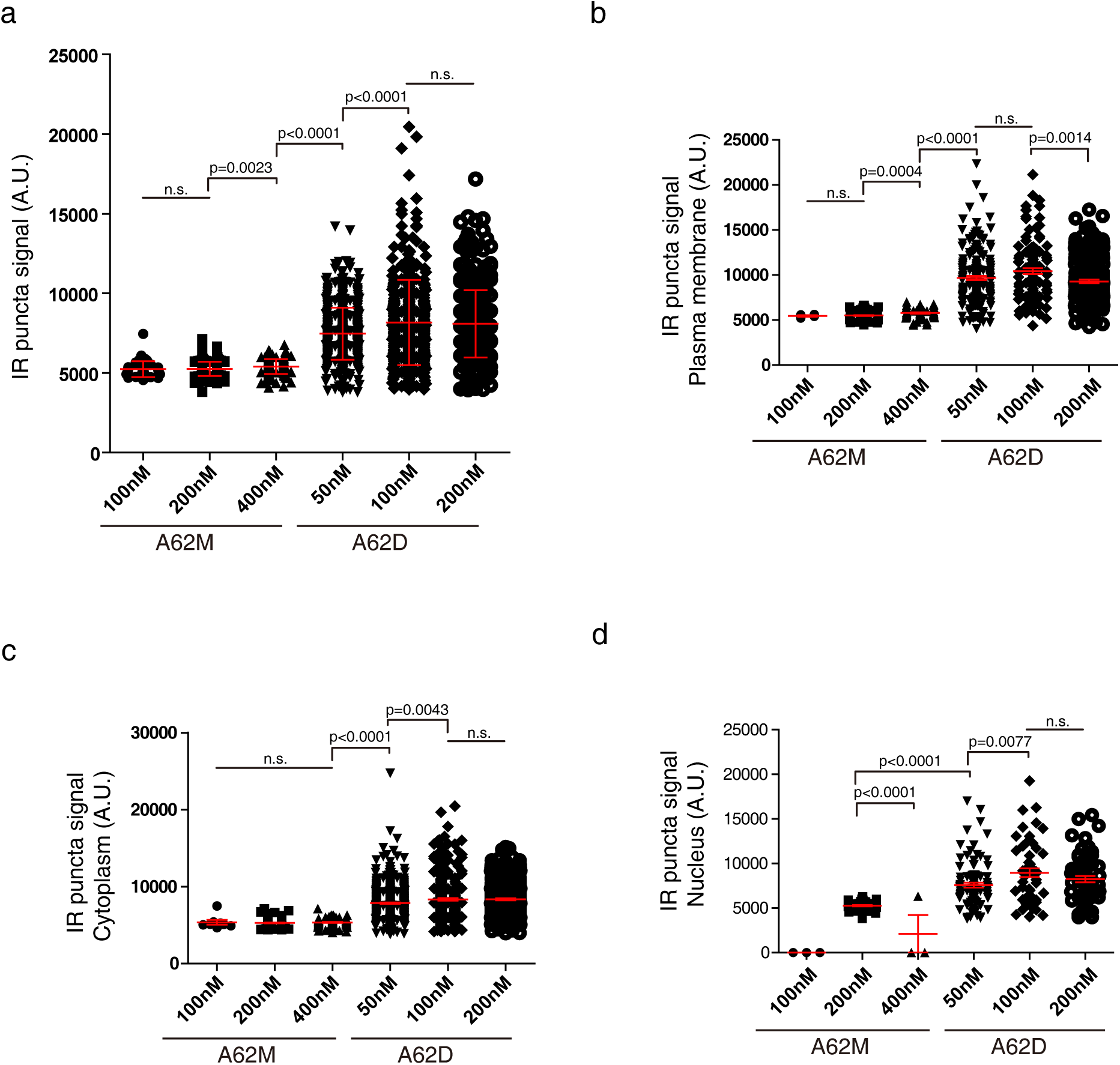
A62D aptamer induces oligomerization of IR. (a) Quantification of IR signal intensities in puncta. Data are presented as mean±SD (standard deviation) to illustrate the distribution of signal intensities. The number of puncta analyzed is as follows: A62M, 100nM: 42; A62M, 200nM: 274; A62M, 400nM: 173; A62D, 50nM: 508; A62D, 100nM: 345; A62D, 200nM: 396. (b) Quantification of IR signal intensities in puncta localized in the plasma membrane. The plasma membrane was defined as the F-actin-stained region, and IR puncta within this region were analyzed. Data are presented as mean±SEM (standard error of the mean). The number of puncta analyzed is as follows: A62M, 100nM: 19; A62M, 200nM: 106; A62M, 400nM: 66; A62D, 50nM: 179; A62D, 100nM: 106; A62D, 200nM: 175. (c) Quantification of IR signal intensities in puncta in the cytoplasm. The cytoplasmic region was defined as the area excluding both the F-actin-and DAPI-stained regions. Data are presented as mean±SEM. The number of puncta analyzed is as follows: A62M, 100nM; 24, A62M, 200nM: 132; A62M, 400nM: 85; A62D, 50nM: 549; A62D, 100nM: 303; A62D, 200nM: 304. (d) Quantification of IR signal intensities in puncta in the nucleus. The nuclear region was defined as the DAPI-stained area. Data are presented as mean±SEM. The number of puncta analyzed is as follows: A62M, 200nM: 38; A62M, 400nM: 3; A62D, 50nM: 94; A62D, 100nM: 52; A62D, 200nM: 60.

**Supplementary Table 1.**
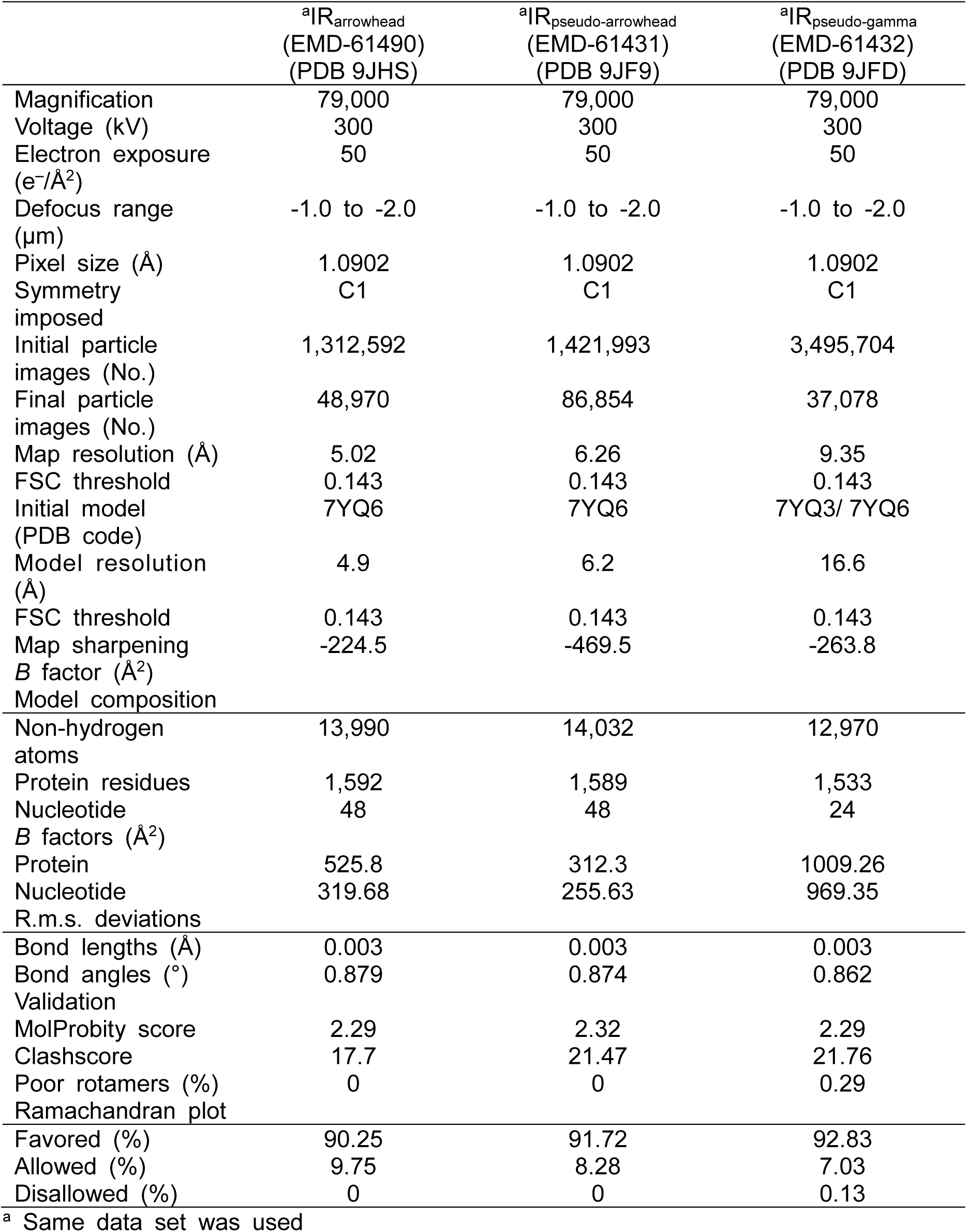
Cryo-EM data collection, refinement and validation statistics.

## References

1. Cohen, D.H. and D. LeRoith, Obesity, type 2 diabetes, and cancer: the insulin and IGF connection. Endocr Relat Cancer, 2012. 19(5): p. F27–45.

2. White, M.F., IRS proteins and the common path to diabetes. Am J Physiol Endocrinol Metab, 2002. 283(3): p. E413–22.

3. Taniguchi, C.M., B. Emanuelli, and C.R. Kahn, Critical nodes in signalling pathways: insights into insulin action. Nat Rev Mol Cell Biol, 2006. 7(2): p. 85–96.

4. Haeusler, R.A., T.E. McGraw, and D. Accili, Biochemical and cellular properties of insulin receptor signalling. Nat Rev Mol Cell Biol, 2018. 19(1): p. 31–44.

5. Lemmon, M.A. and J. Schlessinger, Cell signaling by receptor tyrosine kinases. Cell, 2010. 141(7): p. 1117–34.

6. American Diabetes, A., Diagnosis and classification of diabetes mellitus. Diabetes Care, 2014. 37 Suppl 1: p. S81–90.

7. Gallagher, E.J. and D. LeRoith, Hyperinsulinaemia in cancer. Nat Rev Cancer, 2020. 20(11): p. 629–644.

8. Hubbard, S.R., The insulin receptor: both a prototypical and atypical receptor tyrosine kinase. Cold Spring Harb Perspect Biol, 2013. 5(3): p. a008946.

9. Nielsen, J., et al., Structural Investigations of Full-Length Insulin Receptor Dynamics and Signalling. J Mol Biol, 2022. 434(5): p. 167458.

10. Xiong, X., et al., Symmetric and asymmetric receptor conformation continuum induced by a new insulin. Nat Chem Biol, 2022. 18(5): p. 511–519.

11. Uchikawa, E., et al., Activation mechanism of the insulin receptor revealed by cryo-EM structure of the fully liganded receptor-ligand complex. Elife, 2019. 8.

12. Weis, F., et al., The signalling conformation of the insulin receptor ectodomain. Nat Commun, 2018. 9(1): p. 4420.

13. Scapin, G., et al., Structure of the insulin receptor-insulin complex by single-particle cryo-EM analysis. Nature, 2018. 556(7699): p. 122–125.

14. Yunn, N.O., et al., A stepwise activation model for the insulin receptor. Exp Mol Med, 2023. 55(10): p. 2147–2161.

15. Choi, E. and X.C. Bai, The Activation Mechanism of the Insulin Receptor: A Structural Perspective. Annu Rev Biochem, 2023. 92: p. 247–272.

16. Lawrence, M.C., Understanding insulin and its receptor from their three-dimensional structures. Mol Metab, 2021. 52: p. 101255.

17. Croll, T.I., et al., Higher-Resolution Structure of the Human Insulin Receptor Ectodomain: Multi-Modal Inclusion of the Insert Domain. Structure, 2016. 24(3): p. 469–76.

18. An, W., et al., Activation of the insulin receptor by insulin-like growth factor 2. Nat Commun, 2024. 15(1): p. 2609.

19. Viola, C.M., et al., Structural conservation of insulin/IGF signalling axis at the insulin receptors level in Drosophila and humans. Nat Commun, 2023. 14(1): p. 6271.

20. Wu, M., et al., Functionally selective signaling and broad metabolic benefits by novel insulin receptor partial agonists. Nat Commun, 2022. 13(1): p. 942.

21. Kirk, N.S., et al., Activation of the human insulin receptor by non-insulin-related peptides. Nat Commun, 2022. 13(1): p. 5695.

22. Li, J., et al., Synergistic activation of the insulin receptor via two distinct sites. Nat Struct Mol Biol, 2022. 29(4): p. 357–368.

23. Gutmann, T., et al., Cryo-EM structure of the complete and ligand-saturated insulin receptor ectodomain. J Cell Biol, 2020. 219(1).

24. Kim, J., et al., Functional selectivity of insulin receptor revealed by aptamer-trapped receptor structures. Nat Commun, 2022. 13(1): p. 6500.

25. Li, J., et al., Structural basis of the activation of type 1 insulin-like growth factor receptor. Nat Commun, 2019. 10(1): p. 4567.

26. DeMeyts, P., A.R. Bainco, and J. Roth, Site-site interactions among insulin receptors. Characterization of the negative cooperativity. J Biol Chem, 1976. 251(7): p. 1877–88.

27. de Meyts, P., et al., Insulin interactions with its receptors: experimental evidence for negative cooperativity. Biochem Biophys Res Commun, 1973. 55(1): p. 154–61.

28. Lawrence, C.F., et al., Insulin Mimetic Peptide Disrupts the Primary Binding Site of the Insulin Receptor. J Biol Chem, 2016. 291(30): p. 15473–81.

29. Hinke, S.A., et al., Unique pharmacology of a novel allosteric agonist/sensitizer insulin receptor monoclonal antibody. Mol Metab, 2018. 10: p. 87–99.

30. Corbin, J.A., et al., Improved glucose metabolism in vitro and in vivo by an allosteric monoclonal antibody that increases insulin receptor binding affinity. PLoS One, 2014. 9(2): p. e88684.

31. Yunn, N.O., et al., An aptamer agonist of the insulin receptor acts as a positive or negative allosteric modulator, depending on its concentration. Exp Mol Med, 2022. 54(4): p. 531–541.

32. Yunn, N.O., et al., Agonistic aptamer to the insulin receptor leads to biased signaling and functional selectivity through allosteric modulation. Nucleic Acids Res, 2015. 43(16): p. 7688–701.

33. Punjani, A., et al., cryoSPARC: algorithms for rapid unsupervised cryo-EM structure determination. Nat Methods, 2017. 14(3): p. 290–296.

34. Bepler, T., et al., Positive-unlabeled convolutional neural networks for particle picking in cryo-electron micrographs. Nat Methods, 2019. 16(11): p. 1153–1160.

35. Pettersen, E.F., et al., UCSF Chimera--a visualization system for exploratory research and analysis. J Comput Chem, 2004. 25(13): p. 1605–12.

36. Emsley, P. and K. Cowtan, Coot: model-building tools for molecular graphics. Acta Crystallogr D Biol Crystallogr, 2004. 60(Pt 12 Pt 1): p. 2126–32.

37. Liebschner, D., et al., Macromolecular structure determination using X-rays, neutrons and electrons: recent developments in Phenix. Acta Crystallogr D Struct Biol, 2019. 75(Pt 10): p. 861–877.

38. Chen, V.B., et al., MolProbity: all-atom structure validation for macromolecular crystallography. Acta Crystallogr D Biol Crystallogr, 2010. 66(Pt 1): p. 12–21.

39. Choi, E. and H. Lee, Chromosome damage in mitosis induces BubR1 activation and prometaphase arrest. FEBS Lett, 2008. 582(12): p. 1700–6.

40. Choi, E., et al., BubR1 acetylation at prometaphase is required for modulating APC/C activity and timing of mitosis. EMBO J, 2009. 28(14): p. 2077–89.

41. Choi, E., et al., BRCA2 fine-tunes the spindle assembly checkpoint through reinforcement of BubR1 acetylation. Dev Cell, 2012. 22(2): p. 295–308.

42. Park, I., et al., Loss of BubR1 acetylation causes defects in spindle assembly checkpoint signaling and promotes tumor formation. J Cell Biol, 2013. 202(2): p. 295–309.

43. Chen, X., et al., Superresolution structured illumination microscopy reconstruction algorithms: a review. Light Sci Appl, 2023. 12(1): p. 172.

44. Riccardi, C., et al., Dimeric and Multimeric DNA Aptamers for Highly Effective Protein Recognition. Molecules, 2020. 25(22).

45. Bujotzek, A., et al., Towards a rational spacer design for bivalent inhibition of estrogen receptor. J Comput Aided Mol Des, 2011. 25(3): p. 253–62.

46. Wang, Z., et al., Multivalent Aptamer Approach: Designs, Strategies, and Applications. Micromachines (Basel), 2022. 13(3).

47. Yunn, N.O., et al., A hotspot for enhancing insulin receptor activation revealed by a conformation-specific allosteric aptamer. Nucleic Acids Res, 2021. 49(2): p. 700–712.

48. Li, H., et al., Mechanism of INSR clustering with insulin activation and resistance revealed by super-resolution imaging. Nanoscale, 2022. 14(20): p. 7747–7755.

49. Dall’Agnese, A., et al., The dynamic clustering of insulin receptor underlies its signaling and is disrupted in insulin resistance. Nat Commun, 2022. 13(1): p. 7522.

50. Spratt, J., et al., Multivalent insulin receptor activation using insulin-DNA origami nanostructures. Nat Nanotechnol, 2024. 19(2): p. 237–245.

51. Bhaskar, V., et al., A fully human, allosteric monoclonal antibody that activates the insulin receptor and improves glycemic control. Diabetes, 2012. 61(5): p. 1263–71.

52. Jensen, M., et al., Activation of the insulin receptor by insulin and a synthetic peptide leads to divergent metabolic and mitogenic signaling and responses. J Biol Chem, 2007. 282(48): p. 35179–86.

53. Park, J., et al., Activation of the insulin receptor by an insulin mimetic peptide. Nat Commun, 2022. 13(1): p. 5594.

54. Xiong, X., et al., A structurally minimized yet fully active insulin based on cone-snail venom insulin principles. Nat Struct Mol Biol, 2020. 27(7): p. 615–624.

55. Menting, J.G., et al., How insulin engages its primary binding site on the insulin receptor. Nature, 2013. 493(7431): p. 241–5.

