## Supplemental Data 1 for "Structural mechanism of insulin receptor activation by a dimeric aptamer agonist"

### Supplementary Information

#### Structural comparison of $IR_{pseudo\text{-}gamma}$ with $IR_{pseudo\text{-}arrowhead}$ and $IR_{arrowhead}$

To investigate the structural changes from $IR_{pseudo\text{-}arrowhead}$ to $IR_{pseudo\text{-}gamma}$, we aligned the stalks of protomer A (Supplementary Fig. 7a). In $IR_{pseudo\text{-}gamma}$, the aptamer-free head (L1-CR domain) of protomer A is elevated by 10.2º compared to the aptamer-bound protomer A of $IR_{pseudo\text{-}arrowhead}$ (Q177-L459-V604; L1-L2-FnIII2). Meanwhile, the aptamer-bound protomer B in $IR_{pseudo\text{-}gamma}$ is downshifted by 7.2º relative to protomer B in $IR_{pseudo\text{-}arrowhead}$ (Supplementary Fig. 7b). The L1 domain of protomer A in $IR_{pseudo\text{-}gamma}$ has shifted by 29.5 Å, while in protomer B, it has shifted by 16.4 Å.

Alignment of the L2 domains between $IR_{pseudo\text{-}gamma}$ and $IR_{pseudo\text{-}arrowhead}$ revealed that protomer A of $IR_{pseudo\text{-}gamma}$ adopts a more compact form (Q177-E204-L459; L1-CR-L2, 52º) compared to protomer A of $IR_{pseudo\text{-}arrowhead}$ (78º). In contrast, protomer B of $IR_{pseudo\text{-}gamma}$ maintains a similar conformation to protomer A of $IR_{pseudo\text{-}arrowhead}$ (86º vs. 83º) (Supplementary Fig. 7c, d). The distance between L1 domains in protomer A and B of $IR_{pseudo\text{-}gamma}$ is 43.3 Å and 6.7 Å, respectively. Overall, protomer A undergoes significant changes at two hinge regions (CR-L2 and L2-FnIII1), while protomer B shows relatively smaller changes, particularly in the L2-FnIII1 hinge.

When aligning the stalk of $IR_{pseudo\text{-}gamma}$ with that of $IR_{arrowhead}$, one head shifts upward while the other shifts downward at the L2-FnIII1 hinge (Supplementary Fig. 7e, f). The ligand-free protomer A is uplifted by 19.3º and 15º at Q177-L459-V604 (L1-L2-FnIII2) and E204-L459-V604 (CR-L2-FnIII2), respectively, while the aptamer-bound protomer B is downshifted by 1.1º and 10.6º. The L1 domain of protomer A shifts by 48 Å, while in protomer B, it shifts by 26 Å.

Upon aligning the L2 domains, the aptamer-free protomer A of $IR_{pseudo\text{-}gamma}$ forms a more compact structure (angle reduced from 82º to 52º), while the aptamer-bound protomer B retains a nearly identical conformation (angle shift from 82º to 86º) (Supplementary Fig. 7g, h). The distance between L1-L1' or CR-CR' is much smaller for protomer B (5 Å) compared to protomer A (60 Å and 44 Å, respectively). Thus, protomer A experiences substantial shifts at two hinges, while the L1-FnIII1 hinge of aptamer-bound protomer B undergoes only minor adjustments.

#### Supplementary Figure Legends

**Supplementary Figure 1. Effects of the dose and duration of A62M and A62D-8T on IR phosphorylation and downstream signaling**
(a) Comparison of dose-dependent phosphorylation of IR and downstream signaling by A62M or A62D-8T. Rat-1/hIR cells were incubated with varying concentrations of A62M or A62D for 1 h. (b) The relative band intensities, compared to 100 nM insulin, are shown for pIR Y1150, pIR Y1150/Y1151, pAKT T308, pAKT S473, and pERK T202/Y204. Quantified values for the band intensities are shown in Fig 2.

**Supplementary Figure 2. Purification of the IR + A62D complex, analysis of the quality of the cryo-EM map** (a) Size-exclusion chromatography profile and SDS-PAGE result. (b) Representative cryo-EM micrograph of the IR + A62D complex. **c-e,** 2D class averages of (c) $IR_{arrowhead}$, (d) $IR_{pseudo\text{-}arrowhead}$ and (e) $IR_{pseudo\text{-}gamma}$. **f-n**, Fourier shell correlation curves (**f-h**), Angular distributions (**i-k**) and Local resolution of the map (**l-n**) of (**f, i, l**) $IR_{arrowhead}$, (**g, j, m**) $IR_{pseudo\text{-}arrowhead}$ and (**h, k, n**) $IR_{pseudo\text{-}gamma}$.

**Supplementary Figure 3. Workflow of cryo-EM processing for the $IR_{arrowhead}$.** Flow chart of data processing for the $IR_{arrowhead}$.

**Supplementary Figure 4. Workflow of cryo-EM processing for the $IR_{pseudo\text{-}arrowhead}$**

**Supplementary Figure 5. Workflow of cryo-EM processing for the $IR_{pseudo\text{-}gamma}$**

**Supplementary Figure 6. Close-up view of the aptamer binding site in $IR_{pseudo\text{-}}$**

**arrowhead** (a) A cartoon representation of the rigid-body rotation of the $IR_{apo}$ (gray) to the $IR_{arrowhead}$ (orange). Only one protomer is shown for clarity. (b) Rigid-body rotation in a is shown in a 90°-rotated view. (c) A close up view of the cryo-EM density for the A62D in the $IR_{pseudo\text{-}arrowhead}$. Only one A62M module of the A62D can be modeled into the density. (d) Cryo-EM density for the A62D in $IR_{pseudo\text{-}arrowhead}$. (e) Interaction of A62M with the L1 and FnIII-1 domains at site l and site u in $IR_{pseudo\text{-}arrowhead}$. (f) Close-up view of the site-l interface, (g) Close-up view of the site-u interface. (h) Superimposed structures of the $IR_{arrowhead}$ and $IR_{pseudo\text{-}arrowhead}$ by aligning protomer A. Comparison of the B protomers between $IR_{arrowhead}$ and $IR_{pseudo\text{-}arrowhead}$ illustrates the rigid-body rotation of protomer B. (i) Aligned structures of $IR_{arrowhead}$ (white) and $IR_{pseudo\text{-}arrowhead}$ (yellow, blue). Distance between the membrane-proximal ends of the FnIII-3 stalks (D907-D907') is shown for both IRs.

**Supplementary Figure 7. Structural comparison of $IR_{pseudo\text{-}gamma}$ with $IR_{pseudo\text{-}arrowhead}$ and $IR_{arrowhead}$**

(a, b) Superimposed structures of protomer A (a) and protomer B (b) in $IR_{pseudo\text{-}gamma}$ (orange) with the respective protomers in $IR_{pseudo\text{-}arrowhead}$ (blue), aligned by stalks. (c, d) Superimposed structures of protomer A (c) and protomer B (d) in $IR_{pseudo\text{-}gamma}$ (orange) with the respective protomers in $IR_{pseudo\text{-}arrowhead}$ (blue) aligned by the L2 domains. Protomers of the $IR_{pseudo\text{-}gamma}$ and $IR_{pseudo\text{-}arrowhead}$ are shown in orange and blue, respectively. (e, f) Superimposed structures of protomer A (e) and protomer B (f) in $IR_{pseudo\text{-}gamma}$ (orange) with the respective protomers in $IR_{arrowhead}$ (blue), aligned by stalks. (g, h) Superimposed structures of protomer A (g) and protomer B (h) in $IR_{pseudo\text{-}gamma}$ with the respective protomers in $IR_{arrowhead}$, aligned by the L2 domains. Protomers of the $IR_{pseudo\text{-}gamma}$ and $IR_{arrowhead}$ are shown in orange and blue, respectively.

**Supplementary Figure 8. A62D aptamer induces oligomerization of IR**

(a) Quantification of IR signal intensities in puncta. Data are presented as mean±SD (standard deviation) to illustrate the distribution of signal intensities. The number of puncta analyzed is as follows: A62M, 100nM: 42; A62M, 200nM: 274; A62M, 400nM:

173; A62D, 50nM: 508; A62D, 100nM: 345; A62D, 200nM: 396.

(b) Quantification of IR signal intensities in puncta localized in the plasma membrane. The plasma membrane was defined as the F-actin-stained region, and IR puncta within this region were analyzed. Data are presented as mean±SEM (standard error of the mean). The number of puncta analyzed is as follows: A62M, 100nM: 19; A62M, 200nM: 106; A62M, 400nM: 66; A62D, 50nM: 179; A62D, 100nM: 106; A62D, 200nM: 175.

(c) Quantification of IR signal intensities in puncta in the cytoplasm. The cytoplasmic region was defined as the area excluding both the F-actin- and DAPI-stained regions. Data are presented as mean±SEM. The number of puncta analyzed is as follows: A62M, 100nM; 24, A62M, 200nM: 132; A62M, 400nM: 85; A62D, 50nM: 549; A62D, 100nM: 303; A62D, 200nM: 304.

(d) Quantification of IR signal intensities in puncta in the nucleus. The nuclear region was defined as the DAPI-stained area. Data are presented as mean±SEM. The number of puncta analyzed is as follows: A62M, 200nM: 38; A62M, 400nM: 3; A62D, 50nM: 94; A62D, 100nM: 52; A62D, 200nM: 60.

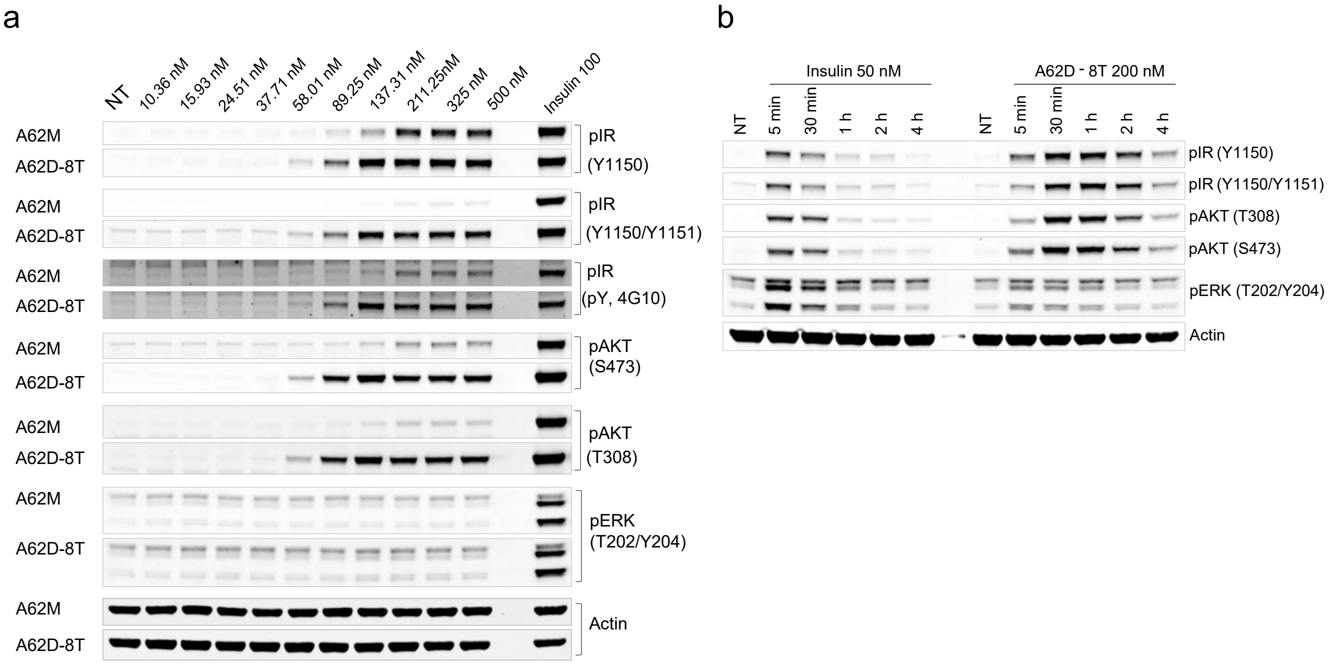


### Supplementary Figure 1

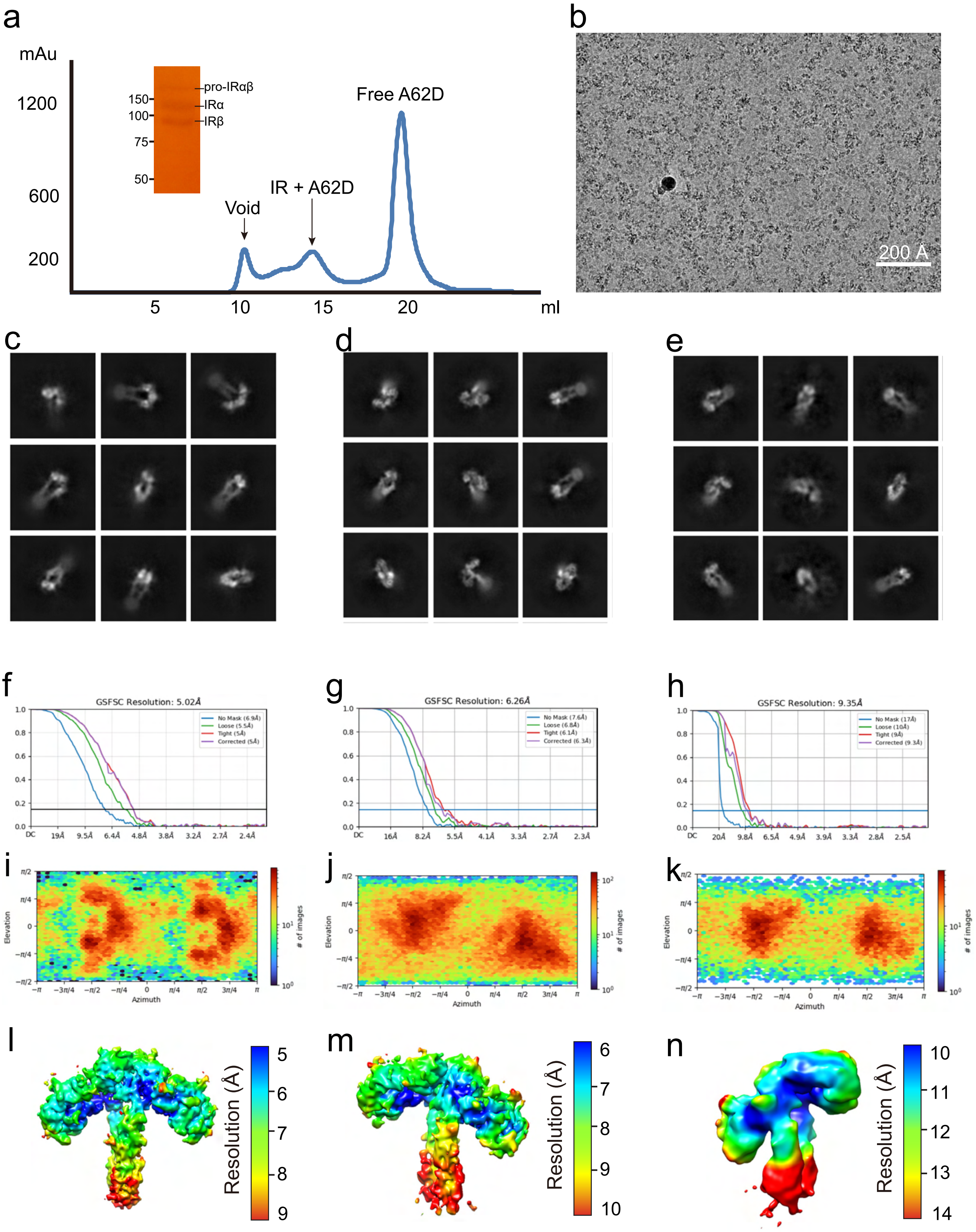


**Supplementary Figure 2**

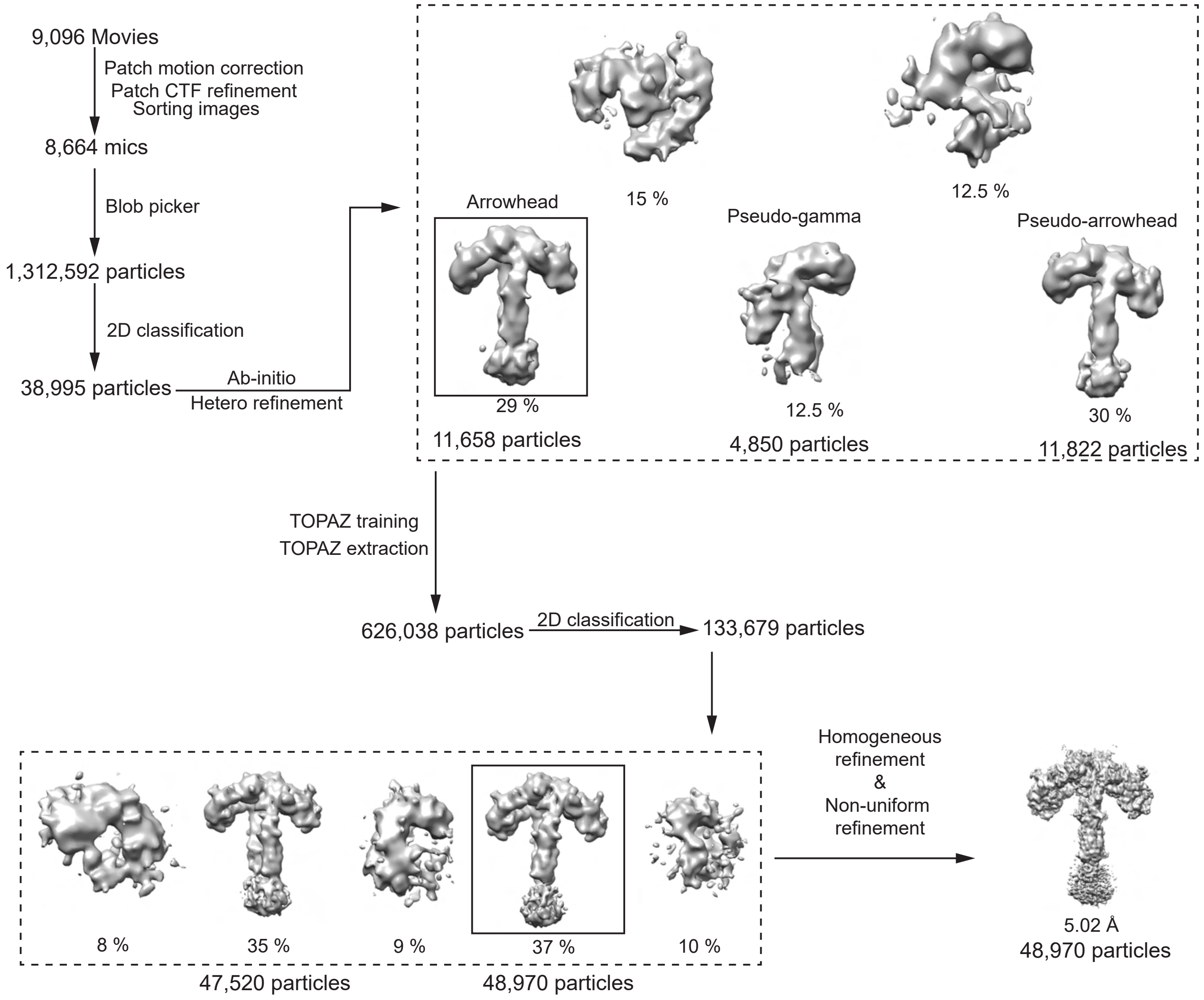


### Supplementary Figure 3

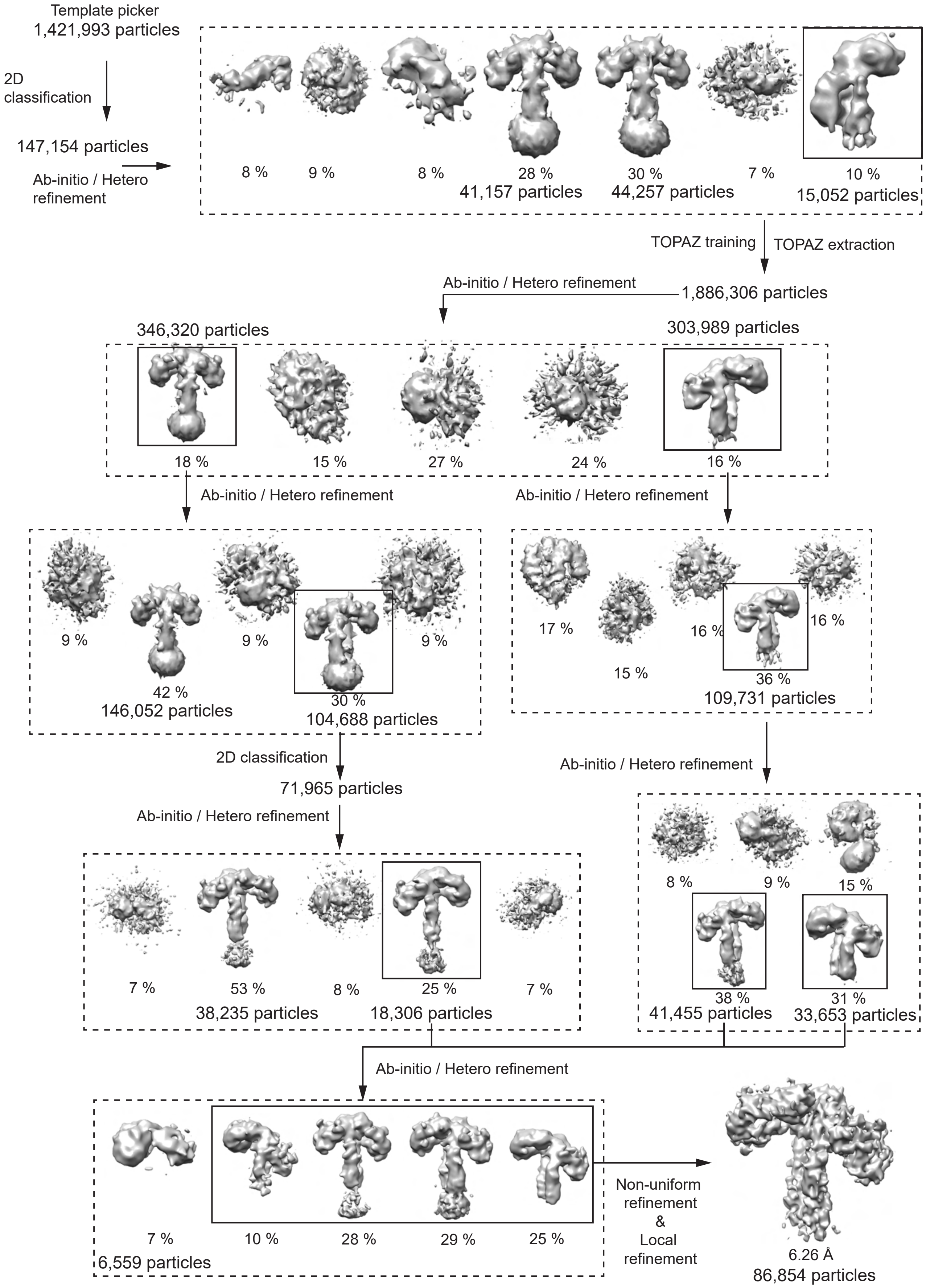


**Supplementary Figure 4**

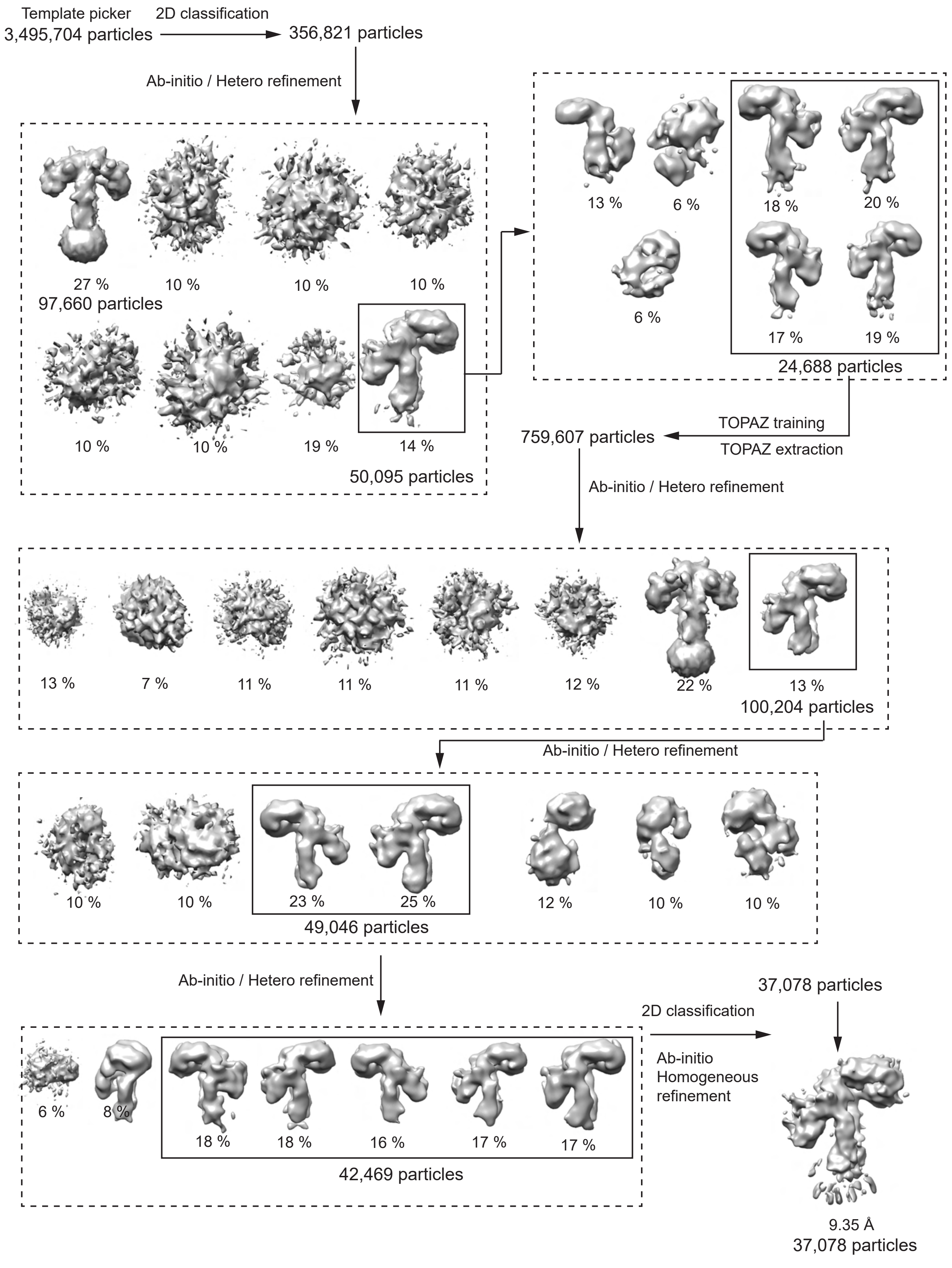


### Supplementary Figure 5

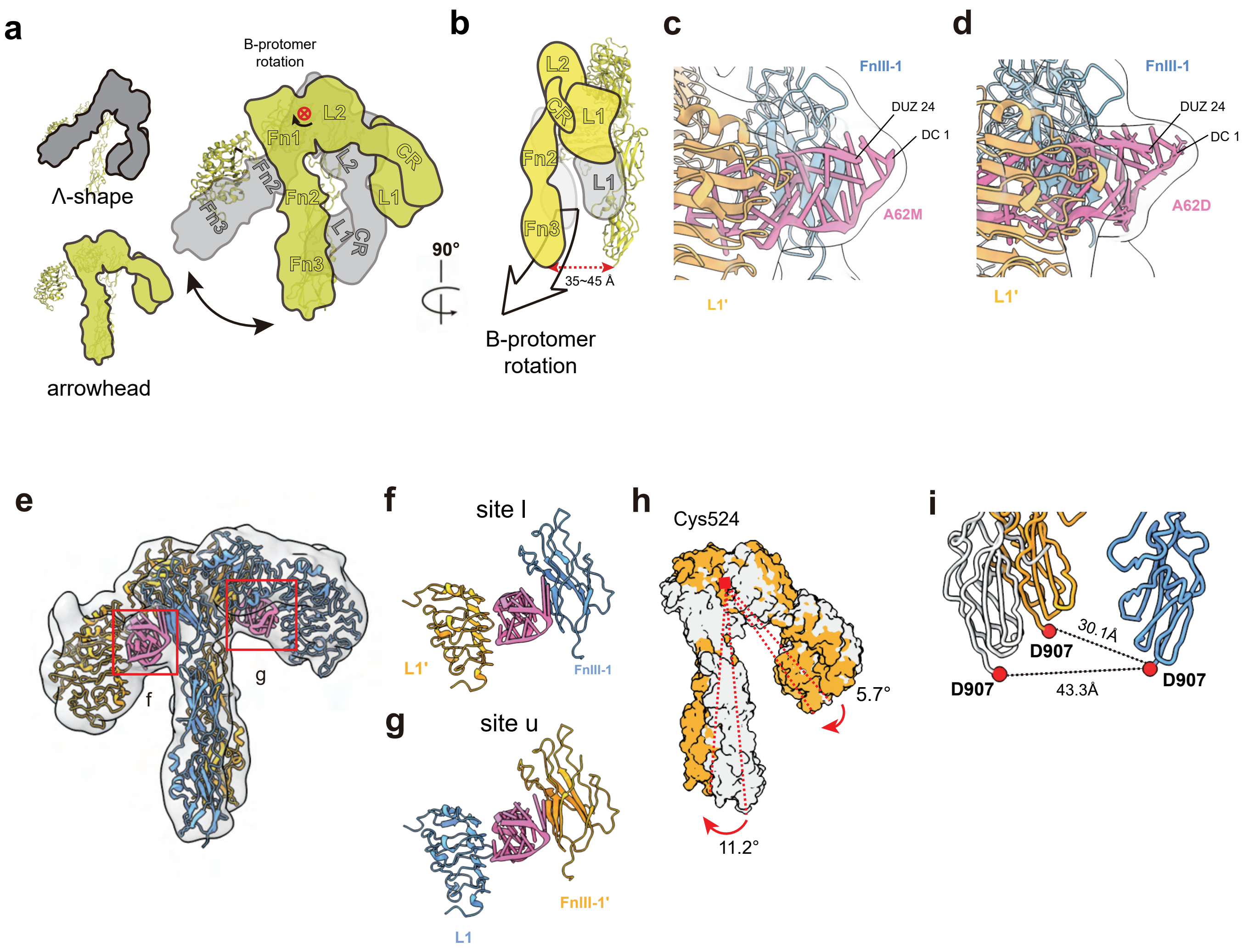


**Supplementary Figure 6**

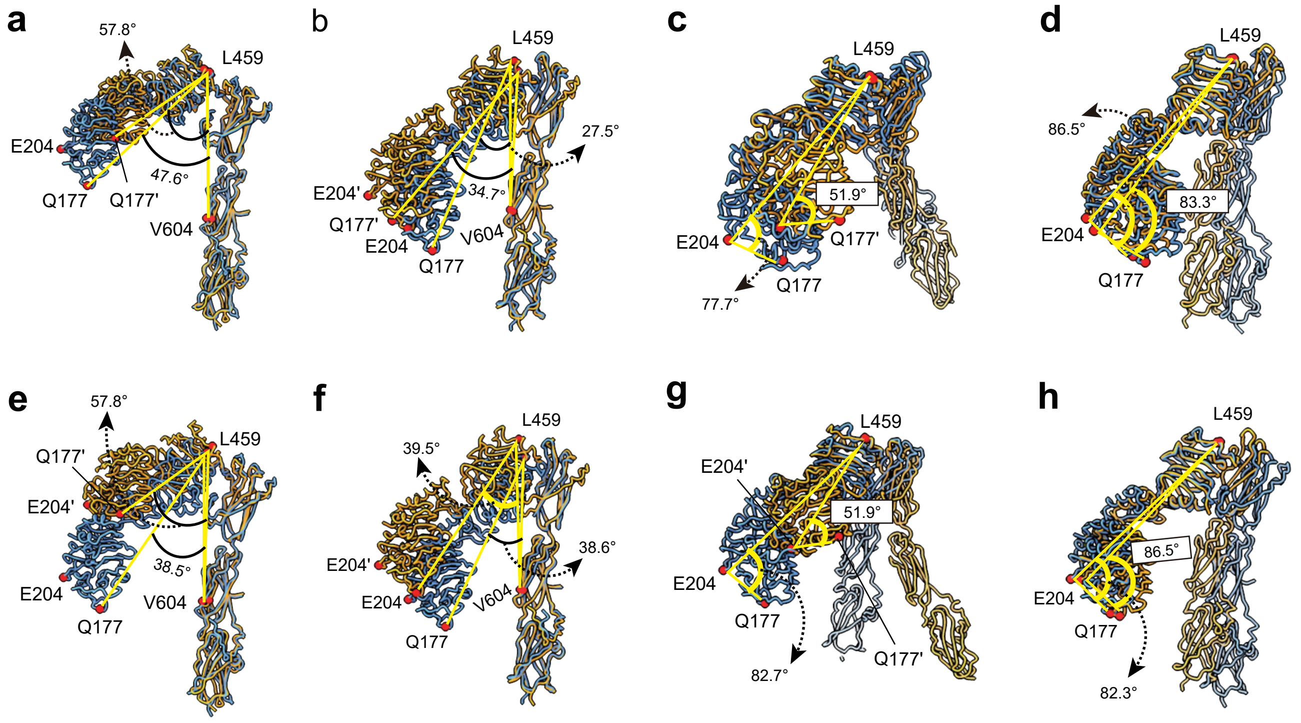


### Supplementary Figure 7

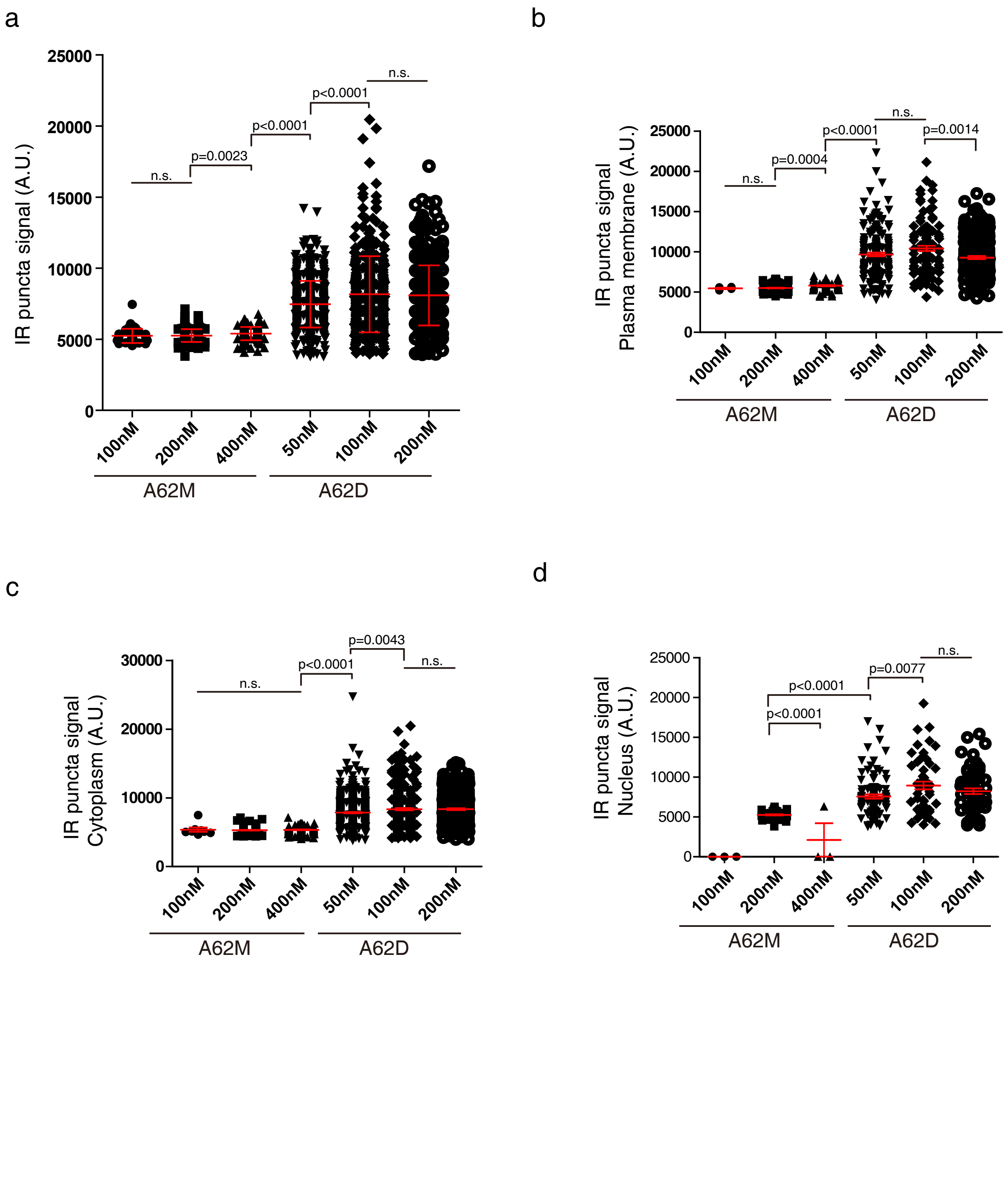


### Supplementary Figure 8

**Supplementary Table 1 | Cryo-EM data collection, refinement and validation statistics.**

| | [a]$IR_{arrowhead}$ (EMD-61490) (PDB 9JHS) | [a]$IR_{pseudo\text{-}arrowhead}$ (EMD-61431) (PDB 9JF9) | [a]$IR_{pseudo\text{-}gamma}$ (EMD-61432) (PDB 9JFD) |
|---|---|---|---|
| Magnification | 79,000 | 79,000 | 79,000 |
| Voltage (kV) | 300 | 300 | 300 |
| Electron exposure ($e^{-}/Å^2$) | 50 | 50 | 50 |
| Defocus range (μm) | -1.0 to -2.0 | -1.0 to -2.0 | -1.0 to -2.0 |
| Pixel size (Å) | 1.0902 | 1.0902 | 1.0902 |
| Symmetry imposed | C1 | C1 | C1 |
| Initial particle images (No.) | 1,312,592 | 1,421,993 | 3,495,704 |
| Final particle images (No.) | 48,970 | 86,854 | 37,078 |
| Map resolution (Å) | 5.02 | 6.26 | 9.35 |
| FSC threshold | 0.143 | 0.143 | 0.143 |
| Initial model (PDB code) | 7YQ6 | 7YQ6 | 7YQ3/ 7YQ6 |
| Model resolution (Å) | 4.9 | 6.2 | 16.6 |
| FSC threshold | 0.143 | 0.143 | 0.143 |
| Map sharpening *B* factor ($Å^2$) | -224.5 | -469.5 | -263.8 |
| Model composition | | | |
| Non-hydrogen atoms | 13,990 | 14,032 | 12,970 |
| Protein residues | 1,592 | 1,589 | 1,533 |
| Nucleotide | 48 | 48 | 24 |
| *B* factors ($Å^2$) | | | |
| Protein | 525.8 | 312.3 | 1009.26 |
| Nucleotide | 319.68 | 255.63 | 969.35 |
| R.m.s. deviations | | | |
| Bond lengths (Å) | 0.003 | 0.003 | 0.003 |
| Bond angles (°) | 0.879 | 0.874 | 0.862 |
| Validation | | | |
| MolProbity score | 2.29 | 2.32 | 2.29 |
| Clashscore | 17.7 | 21.47 | 21.76 |
| Poor rotamers (%) | 0 | 0 | 0.29 |
| Ramachandran plot | | | |
| Favored (%) | 90.25 | 91.72 | 92.83 |
| Allowed (%) | 9.75 | 8.28 | 7.03 |
| Disallowed (%) | 0 | 0 | 0.13 |

[a] Same data set was used
